# Memory out of context: Spacing effects and decontextualization in a computational model of the medial temporal lobe

**DOI:** 10.1101/2022.12.01.518703

**Authors:** James Antony, Xiaonan L. Liu, Yicong Zheng, Charan Ranganath, Randall C. O’Reilly

## Abstract

Some neural representations change across multiple timescales. Here we argue that modeling this “drift” could help explain the spacing effect (the long-term benefit of distributed learning), whereby differences between stored and current temporal context activity patterns produce greater error-driven learning. We trained a neurobiologically realistic model of the entorhinal cortex and hippocampus to learn paired associates alongside temporal context vectors that drifted between learning episodes and/or before final retention intervals. In line with spacing effects, greater drift led to better model recall after longer retention intervals. Dissecting model mechanisms revealed that greater drift increased error-driven learning, strengthened weights in slower-drifting temporal context neurons (temporal abstraction), and improved direct cue-target associations (decontextualization). Intriguingly, these results suggest that decontextualization — generally ascribed only to the neocortex — can occur within the hippocampus itself. Altogether, our findings provide a mechanistic formalization for established learning concepts such as spacing effects and errors during learning.

## Introduction

A primary goal in learning is to make information accessible long after encoding. One well-known technique for enhancing retention is distributing learning events over time rather than cramming them into a short interval. This phenomenon, known as the spacing effect (Maddox, 2016; Russo, Parkin, Taylor, & Wiks, 1998), is ubiquitous across many memory paradigms (Cepeda, Pashler, Vul, Wixted, & Rohrer, 2006; Russo et al., 1998; C. D. Smith & Scarf, 2017) and operates over a wide spectrum of time scales spanning seconds (Glenberg, 1976), days (Cepeda et al., 2006; Küpper-Tetzel, Kapler, & Wiseheart, 2014), months (Cepeda et al., 2009; Cepeda, Vul, Rohrer, Wixted, & Pashler, 2008), and years (Bahrick, Bahrick, Bahrick, & Bahrick, 1993).

While the spacing effect has profound real-world implications, it also presents a theoretical puzzle: how does the passage of time, which normally leads to forgetting, also allow for dramatically better long-term memory strengthening after further learning? We propose that one part of the answer is that forgetting can actually enable stronger learning the next time, to the extent that learning is based on the *difference* between the existing memory strength and the full memory strength at the time of learning. This type of learning is known as error-driven learning (EDL), which plays a central role in our model. The other key element to this puzzle involves the role of *context* in both memory encoding and recall. Decades of research has shown that spatial, temporal, situational, and mental contexts contribute to the what-when-where of episodic memories for everyday learning events (Davachi, 2006; Eichenbaum, Yonelinas, & Ranganath, 2007). Each of these contextual factors can support holistic episodic memory recall when cued later, and they are each represented within the intricate neural machinery of the hippocampus (HC) and its major input, the entorhinal cortex (EC). However, just as contextual cues support memory, we will argue that they also limit the potential use of memories to the instances in which they can be reinstated. In order for learning to be accessible over longer intervals, memories may benefit from becoming temporally abstracted or “decontextualized” – two ways that the memory can generalize beyond the local, learned context (Karpicke, Lehman, & Aue, 2014; S. M. Smith & Handy, 2014). As explained further below, these processes constitute a major part of how our computational model produces spacing effects.

Among the many dimensions of context, the most relevant for the spacing effect is temporal context. A collection of computational models have posited that, as memories are encoded, temporal context can be represented as a distributed, drifting pattern of neural activity (Balota, Duchek, & Paullin, 1989; Estes, 1955a, 1955b; Horner, Bisby, Wang, Bogus, & Burgess, 2016; Howard & Kahana, 2002; Howard, Youker, & Venkatadass, 2008; Kahana, 2020; Kiliç, Criss, & Howard, 2013; L. J. Lohnas, Polyn, & Kahana, 2015; Mensink & Raaijmakers, 1988; Mozer, Pashler, Cepeda, Lindsey, & Vul, 2009; Murdock, 1997; Polyn, Norman, & Kahana, 2009; Raaijmakers, 2003; Rouhani, Norman, Niv, & Bornstein, 2020; Sederberg, Gershman, Polyn, & Norman, 2011). One consequence of this arrangement is that idiosyncratic, encoding-related activity patterns become reinstated during retrieval (El-Kalliny et al., 2019; Folkerts, Rutishauser, & Howard, 2018; Howard, Viskontas, Shankar, & Fried, 2012; Manning, Polyn, Baltuch, Litt, & Kahana, 2011). In our model, this drifting temporal context provides a well-established explanation for the temporal forgetting function, in terms of the gradually diminishing contextual cue support for the memory (Bouton, 1993; Crowder, 1976; Estes, 1955a; Gershman & Niv, 2010; Mensink & Raaijmakers, 1988). Moreover, greater drift creates greater mismatches between the temporal contexts at encoding and re-learning, enhancing the EDL that then drives greater plasticity for more widely spaced items. In addition, we show that this form of learning favors the elements in common between the two learning events, resulting in better subsequent recall that relies less on the temporal context. Thus, our model demonstrates how spacing effects emerge as a synergistic interaction between contextual drift and EDL (Mozer et al., 2009).

In the following introductory sections, we first discuss prior spacing effect findings and theories. Second, we provide behavioral and neural evidence that temporal context drifts across multiple time scales. Third, we tie these various lines of evidence together to introduce a drifting, biologically plausible model of the entorhinal cortex and hippocampus to simulate paired associate learning. Finally, we discuss how and what gets strengthened due to spacing, how this compares against prior spacing effect theories, and what this means for the fate of the memory.

### The non-monotonic relationship between spacing and final retention interval and its explanation

Spacing effects have received considerable attention in the cognitive psychology literature, generating a rich array of findings and explanations. Typically, longer intervals between learning events (inter-stimulus intervals, or ISIs) produce superior memory after some retention interval (RI) following the last instance of learning. However, one curious result is that more spacing is not always better. Rather, the optimal ISI depends strongly on RI, such that for very short RIs, short ISIs (or “massed” trials) often confer an advantage over spacing (Balota et al., 1989; Glenberg, 1976; Peterson, Wampler, Kirkpatrick, & Saltzman, 1963; Rawson & Kintsch, 2005; Spieler & Balota, 1996; Toppino & Gerbier, 2014). In fact, plots relating ISI to RI are often non-monotonic, with recall (given the same RI) rising sharply, reaching a maximum, and slowly decreasing with increasing ISI (Cepeda et al., 2009, 2006, 2008). No single ISI always benefits memory the most, and therefore, memory performance cannot be explained by a single factor – the relationship is more complex.

One prominent explanation for spacing effects is encoding variability theory. This theory suggests that a greater temporal difference between learning events results in more temporal unique elements assigned to each learning instance of the memory. This creates more variable encoding contexts during learning that in turn allow for more routes to the memory during retrieval (Balota, Duchek, & Logan, 2007; Glenberg, 1976, 1979; Huff & Bodner, 2014; L. Lohnas, Polyn, & Kahana, 2011; McFarland, Rhodes, & Frey, 1979; Melton, 1970; Raaijmakers, 2003; Ross & Landauer, 1978). However, as discussed above, maximum spacing (and hence, maximum encoding variability) does not always produce maximum memory benefits, making encoding variability alone unsatisfying as an explanation of spacing. Therefore, encoding variability has often been paired with another process called study-phase retrieval (Benjamin & Tullis, 2010; Cepeda et al., 2009; Greene, 1989; Mozer et al., 2009; Raaijmakers, 2003). Study-phase retrieval suggests that memory strengthening only occurs if elements of the study phase can actually be retrieved and reactivated during relearning (Thios & D’Agostino, 1976). Given that this ability will decrease over time, there will be a lower likelihood of strengthening at later intervals. On its own, study-phase retrieval would produce an *anti-spacing* effect, but combining it with the benefits afforded by encoding variability offers a plausible account of how one might observe non-monotonic effects between ISI and RI in the spacing effect (Benjamin & Tullis, 2010; Landauer, 1969; Maddox, 2016; Raaijmakers, 2003).

While encoding variability is plausible in explaining many of the behavioral effects of spacing, the spacing effect is a temporal phenomenon, and encoding variability largely does not incorporate recent developments on the critical role of temporal context for memory [though see Raaijmakers (2003)]. Here, we will argue that the non-monotonic nature of spacing effects can be explained via EDL as a strengthening mechanism. We propose that adding *more* routes to a memory trace, which is the memory-strengthening mechanism proposed by encoding variability theory, may be less important than *strengthening* aspects of the trace in common across learning events.

### Behavioral and neural evidence of multi-scale drift in temporal context

A central feature of episodic memory is that it is temporally organized. One classic example demonstrating this organization is that, during free recall, subjects tend to recall information successively that was presented nearby during learning (Antony et al., 2021; Healey, 2018; Healey, Long, & Kahana, 2019; Heusser, Ezzyat, Shiff, & Davachi, 2018; Howard & Kahana, 2002; Kahana, 1996, 2020; Uitvlugt & Healey, 2019). Explanations of this effect center around the temporal context model (Howard & Kahana, 2002; L. J. Lohnas et al., 2015; Polyn et al., 2009; Sederberg, Howard, & Kahana, 2008), which shows that a slowly drifting temporal context representation at encoding can become reinstated with a corresponding episodic memory, guiding subsequent memory recall to other information learned contiguously in time. Importantly, evidence for behavioral temporal contiguity spans numerous time scales. For instance, Howard et al. (2008) had subjects learn a number of word lists and, after the final list, they were asked to recall all words from all lists. They found that subjects did not only transition to nearby words during recall, but also to nearby lists, suggesting temporal context has the property of being scale-invariant. Evidence for scale-invariant temporal representations has also arisen in behavioral paradigms like temporal estimation (Gibbon, 1977; Gibbon, Church, & Meck, 1984; Lewis & Miall, 2009; Merchant, Harrington, & Meck, 2013), other memory paradigms, (Brown, Neath, & Chater, 2007; Singh, Oliva, & Howard, 2017) and a variety of neural data (Bright et al., 2020; Cao, Bladon, Charczynski, Hasselmo, & Howard, 2022; Folkerts et al., 2018; Guo, Huson, Macosko, & Regehr, 2021; Jeunehomme & D’Argembeau, 2020; Manning et al., 2011; Nielson, Smith, Sreekumar, Dennis, & Sederberg, 2015; Rossi-Pool et al., 2019; Yaffe et al., 2014). Accordingly, spectral temporal representations have been incorporated into a variety of computational models of time (Brown et al., 2007; Grossberg & Schmajuk, 1989; Howard, 2018; Howard & Kahana, 2002; Jacques, Tiganj, Sarkar, Howard, & Sederberg, 2021; Lewandowsky, Ecker, Farrell, & Brown, 2012; Liu, Tiganj, Hasselmo, & Howard, 2019; Miall, 1989; Rolls & Mills, 2019; Tiganj, Hasselmo, & Howard, 2015), and the organization of time has been related to other laws of scale-invariance such as the Weber-Fechner law of perception (Arzy, Adi-Japha, & Blanke, 2009; Brietzke & Meyer, 2021; Cao et al., 2022; Dehaene, 2003). Intriguingly, scale-invariant temporal representations could also explain the shape of forgetting: if temporal context provides cue support for memories, drift over a spectrum of time scales would produce forgetting curves resembling human episodic memory data – that is, forgetting would proceed rapidly, and then more slowly, like a canonical forgetting curve (Ebbinghaus, 1885; D. C. Rubin & Wenzel, 1996).

Scale-invariant temporal context theories of memory have received support from recent neurobiological studies of the EC and HC. EC neurons drift, or slowly increase or decrease in activity, at varying rates over time, from seconds to hours (Aghajan, Kreiman, & Fried, 2022; Bright et al., 2020; Tsao et al., 2018; Umbach et al., 2020). For example, Tsao et al. examined lateral EC (LEC) neurons in rats as they repeatedly explored two different environments over the course of an hour. Activity in a large proportion of recorded neurons drifted at varying rates, including some that drifted slowly over the entire session. Such an arrangement suggests the full population vector of LEC neurons drifts in a multi-scale fashion over time. Within HC, representational drift of neuronal ensembles has been demonstrated over even wider scales, from seconds to months (Devalle & Roxin, 2022; Geva, Deitch, Rubin, & Ziv, 2023; Hainmueller & Bartos, 2018; Keinath & Brandon, 2022; Kinsky, Sullivan, Mau, Hasselmo, & Eichenbaum, 2018; J. S. Lee, Briguglio, Cohen, Romani, & Lee, 2020; Liu et al., 2022; Mankin, Diehl, Sparks, Leutgeb, & Leutgeb, 2015; Manns, Howard, & Eichenbaum, 2007; Mau, Hasselmo, & Cai, 2020; Mau et al., 2018; A. Rubin, Geva, Sheintuch, & Ziv, 2015; Umbach, Tan, Jacobs, Pfeiffer, & Lega, 2022; Y. Ziv et al., 2013), and distinct memories acquired within short temporal windows share greater HC representational overlap (Cai et al., 2016; Rashid et al., 2016; Shen et al., 2022). Additionally, HC in rodents supports temporal order memory (Dusek & Eichenbaum, 1997; Fortin, Agster, & Eichenbaum, 2002) and has cells that chart out repeated intervals of time, or “time cells” (Liu et al., 2019; MacDonald, Lepage, Eden, & Eichenbaum, 2011; Pastalkova, Itskov, Amarasingham, & Buzsáki, 2008; Reddy et al., 2021; Shimbo, Izawa, & Fujisawa, 2021). Evidence from human fMRI experiments also suggests EC and HC support temporal representations. EC may support judgments of temporal duration (Lositsky et al., 2016), and anterolateral EC (alEC) (the analogue of LEC that can be measured in human fMRI) may aid in recalling the temporal context of a movie (Montchal, Reagh, & Yassa, 2019) and representing the temporal proximity of experience during learning (Bellmund, Deuker, & Doeller, 2019). Additionally, HC in fMRI is sensitive to short temporal durations (Barnett, O’Neil, Watson, & Lee, 2014), temporal proximity (Dimsdale-Zucker et al., 2020; Ezzyat & Davachi, 2014), sequences (Hsieh, Gruber, Jenkins, & Ranganath, 2014), and the time since encoding (Jenkins & Ranganath, 2010; Nielson et al., 2015). Altogether, these results suggest that the neural substrate for representing the temporal context of episodes, as simulated in computational models, may be instantiated in LEC and HC [see also Noulhiane, Pouthas, Hasboun, Baulac, and Samson (2007)].

### Present work: A drifting model of the entorhinal cortex and hippocampus

The present modeling work investigates forgetting and spacing effects within a biologically realistic computational model of the medial temporal lobe, drawing inspiration from computational models of drifting, multi-scale temporal contexts [e.g., Estes (1955a); Howard and Kahana (2002); Mensink and Raaijmakers (1988); Tiganj et al. (2015)] and neurobiological evidence of drift [e.g., Tsao et al. (2018)]. Our model builds on prior complementary learning systems (CLS) models (Ketz, Morkonda, & O’Reilly, 2013; McClelland, McNaughton, & O’Reilly, 1995; K. A. Norman & O’Reilly, 2003; Rudy & O’Reilly, 2001) and includes neural network layers for HC subregions like the dentate gyrus (DG), CA3, and CA1, which play distinct roles in episodic memory (Hasselmo & McGaughy, 2004; Leutgeb, Leutgeb, Treves, Moser, & Moser, 2004; Schapiro, Turk-Browne, Botvinick, & Norman, 2017). The model also includes EC, which provides the main input into HC and receives its outputs (Witter, Doan, Jacobsen, Nilssen, & Ohara, 2017), and we now include multi-scale temporal context inputs to EC, simulating the timing of various learning schedules resembling human experiments. We therefore call it the HipSTeR (Hip-pocampus with Spectral Te-mporal Representations) model. The CLS configuration has multiple advantages in explaining episodic memory: it demonstrates how some episodic memory effects arise via specialized machinery, as in how high inhibition in the DG allows similar patterns to be separated (pattern separation) (Grossberg, 1982; O’Reilly & McClelland, 1994; Rolls, 1989; Wigström, 1973) and how an area with strong intra-connections (like CA3) allows prior patterns to be recovered given incomplete inputs (pattern completion) (Colgin, Moser, & Moser, 2008; Hasselmo & McGaughy, 2004; Marr, 1971; K. A. Norman & O’Reilly, 2003; Rolls & Kesner, 2006; Treves & Rolls, 1994; Whittington et al., 2020); it shows how new learning can avoid catastrophic interference of old information by including contextual information as an input (Masse, Grant, & Freedman, 2018; O’Reilly & Munakata, 2000); and it elucidates how different subregions and pathways of the hippocampus contribute independently to episodic memory effects (Schapiro et al., 2017; Zheng, Liu, Nishiyama, Ranganath, & O’Reilly, 2022). Importantly, the model learns via a balance of associative (or Hebbian) learning and EDL processes (O’Reilly & Munakata, 2000), the latter of which we will show to be particularly relevant for simulating spacing effects.

Previous explanations of spacing effects center around encoding variability, whereas in our model, spacing effects arise largely via EDL. EDL conceptually dates back at least to learning rules created by Rescorla and Wagner (1972) and has been applied to learning & memory domains (Brod, 2021; Ku, Hargreaves, Wirth, & Suzuki, 2021; Metcalfe, 2017) and computational models (Sutton & Barto, 1981; Zheng et al., 2022). Principally, in EDL, network weights that underlie memory traces change proportionally to the difference between network predictions based on activations and stored weights versus actual outcomes driven by current inputs (Ketz et al., 2013; O’Reilly, 1996; Zheng et al., 2022). As a memory strengthening mechanism, encoding variability shares some similarities with the error-based account we posit, in that it explains how greater benefits should accrue with greater spacing. However, the reason for these benefits differ between the two accounts. Encoding variability predicts that spacing produces more contextual routes to retrieval. This largely assumes that the two events are independent at encoding, rather than the later event at least partially updating the memory (Hintzman, 1986; Ross & Landauer, 1978). Note that the independence assumption is problematic due to the spacing property of super-additivity. Super-additivity refers to higher probability of recalling a repeated item than either of two items separated by the same temporal lag (Begg & Green, 1988; Benjamin & Tullis, 2010; Maddox, 2016). That is, recall is above what would be obtained if you accounted for the recall likelihoods of two independent stimuli, indicating that there is no independence – rather, the original trace is reactivated. Conversely, the error-based account predicts first that the two learning instances interact (the later instance reactivates the first) and, more specifically, that greater temporal mismatch will produce stronger weight changes to the earlier trace via EDL. Critically, these greater weight changes will strengthen units in common across the memories. In other words, encoding variability predicts spacing effects arise via *more* connections, whereas EDL predicts they arise via *stronger* connections.

If temporal drift occurs, which elements of a prior memory remain in common during a re-learning event? We discuss two strengthening processes relevant to this idea: temporal abstraction (Toppino & Gerbier, 2014) and decontextualization (S. M. Smith & Handy, 2014). Regarding temporal abstraction, if drift occurs over multiple time scales, greater spacing between training examples will result in relatively more overlap in slow-drifting units than fast-drifting units. Although there will be a high temporal mismatch and therefore greater EDL, the weight changes from fast-drifting neurons will keep strengthening new units because they will have drifted too substantially to strengthen the old ones. However, because slow-drifting units retain greater overlap with the prior training examples, the greater EDL can actually strengthen these prior connections. As a result, greater spacing preferentially improves weight strengths of units with longer time scales. Temporal abstraction fits nicely with rational accounts of spacing effects, which posit that, given computational constraints, it may be beneficial to support a memory in proportion to how long it has been since it was last encountered (Anderson & Milson, 1989; Brea, Urbanczik, & Senn, 2014; Kording, Tenenbaum, & Shadmehr, 2007; Mozer et al., 2009). That is, information repeated numerous times within a short interval (e.g., hourly) and then not again will likely only be relevant for the next few hours, whereas information repeated on broader interval (e.g., monthly) will likely be important for longer, and it would be optimal to strengthen units according to this expectancy.

Decontextualization takes the idea of abstraction one step further, suggesting that memories may become resistant to all contextual drift. This process occurs via direct strengthening between the core elements of a memory itself rather than a preferential strengthening of any manner of temporal context units; in the case of paired associate learning (which will be the focus of our modeling efforts), this involves direct weights between cue and target representations. Behavioral evidence for decontextualization from different kinds of contexts shows that, in comparison to constant learning contexts, varying learning contexts slows down learning but improves final memory performance when tests are given in a new context (Glenberg, 1979; Imundo, Pan, Bjork, & Bjork, 2021; S. M. Smith, Glenberg, & Bjork, 1978; S. M. Smith & Handy, 2014, 2016). In other words, varying learning contexts makes the memory depend less on elements of the original learning context at retrieval. Intriguingly, temporal spacing and environmental context changes additively benefit memories tested days later in a novel context (S. M. Smith, 1982; S. M. Smith & Rothkopf, 1984). However, to our knowledge, decontextualization has not been used as a concept to explain spacing effects, whereby memories become less reliant on cue support from the temporal context for retrieval.

As a result of these strengthening processes, we suggest that “cramming” trials in time (relative to distributing them) produces *overfitting* to a local temporal context. These short ISIs can benefit memories when RIs are also short. However, after substantial temporal drift occurs (like with long RIs), the overfitting from short ISIs ultimately prevents the information from remaining accessible, whereas the temporal abstraction and decontexualization processes that occur with long ISIs keep the memory accessible (Figure 1). Therefore, we will argue that drift is not merely noise, or a nuisance, but may optimize memory function within a computationally constrained system. That is, drift allows old, non-repeated information to be rationally forgotten while strengthening repeated information according to its temporal regularity (Anderson & Milson, 1989; Mozer et al., 2009).

**Figure 1:**
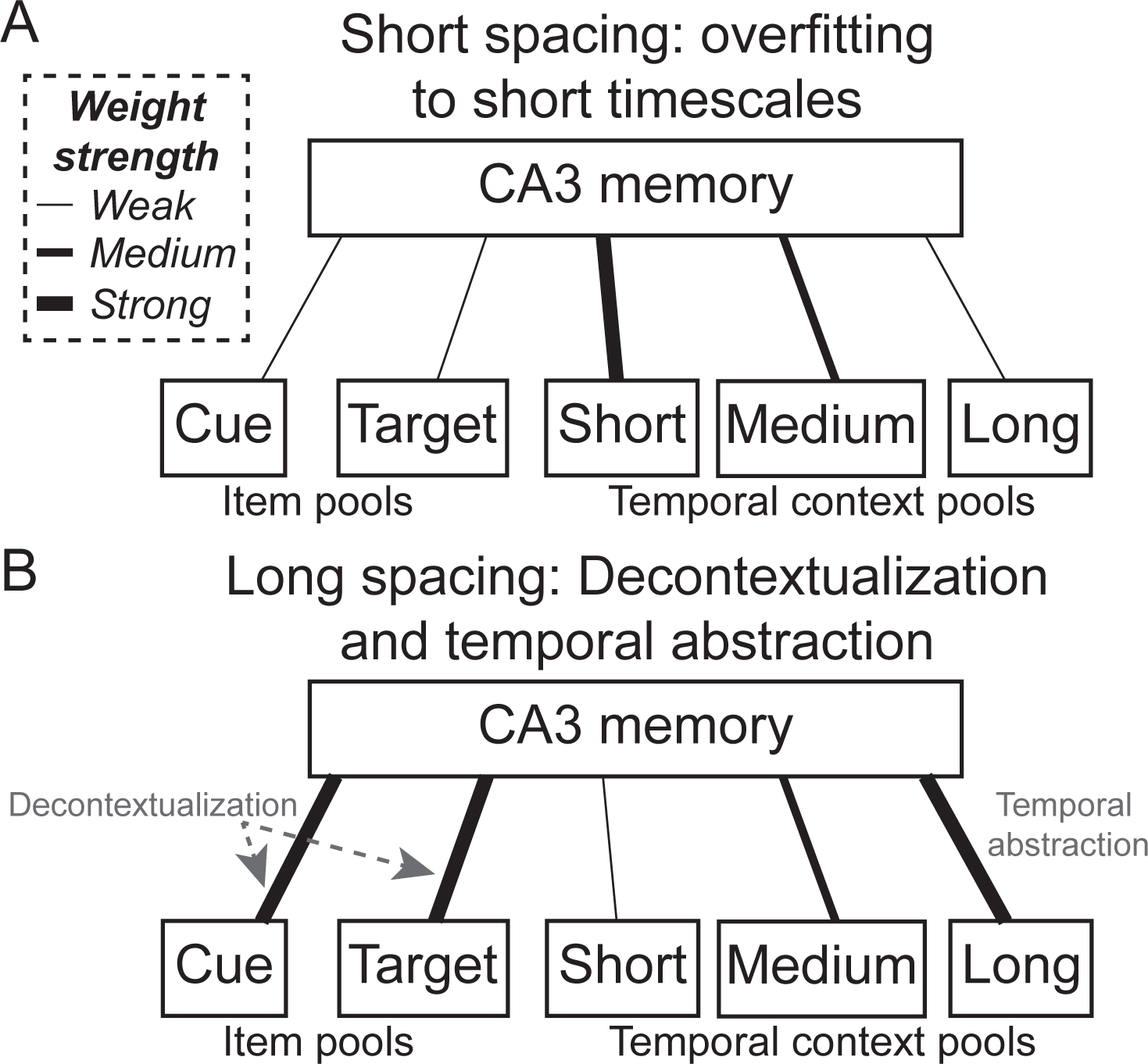
Schematic of basic learning principles under paired associate learning in hippocampal area CA3. (A) Under low spacing, training will preferentially strengthen short timescale representations to CA3, effectively overfitting to the trained temporal context. (B) Under high spacing, greater error-driven learning will be allocated to associations between cues and targets to CA3 (decontextualization, stronger weights on left) and longer timescale representations (temporal abstraction, stronger weights on right). Both processes allow memories to remain accessible after greater drift occurs.

Notably, the error-based model of spacing effects we propose here has an important and related an-tecedent. Mozer et al. (2009) simulated the spacing effect using a multi-scale neural network model, whereby errors in representing temporal context at one timescale of drift were passed to representations at a longer timescale. Alongside the similarities, our approach also differs from theirs in numerous ways: (1) we simulated additional spacing effects, (2) we used a biologically realistic framework of EC-HC, which allowed us to more closely link the network’s learning mechanisms to known neurobiology and create testable neural predictions, (3) temporal abstraction, which was built specifically in Mozer et al. (2009) by passing errors up one layer of the temporal hierarchy, emerged spontaneously in our model without such engineering, and (4) our model suggested decontextualization can also drive spacing benefits. In so doing, our model builds bridges between more abstract models of spacing effects (Mozer et al., 2009; Raaijmakers, 2003; Walsh et al., 2018) and the underlying neural mechanisms in the EC-HC, which opens up more avenues for testing the underlying learning mechanisms and their implications for real-world memory performance.

## Methods

### Model Architecture

Learning in neural networks occurs via the modification of synaptic weights between sending and receiving neurons. Our model was implemented in the Leabra (Local, Error-driven, and Associative, Biologically Realistic Algorithm) framework, which features two distinct learning rules. The first is Hebbian learning, which posits that changes to weights between connected units are incrementally updated through simultaneous, repeated activations (Hebb, 1949). The second, more powerful learning rule is EDL. This rule posits that the network constantly produces expectations (based on activations and stored weights) that are measured against outcomes, and that learning is proportional to the difference between the two (O’Reilly & Munakata, 2000). The model also used rate-coded neurons separated into different layers and pools, sparse and distributed representations, competition driven by inhibitory interneurons within and across layers, and full bidirectional connectivity between some layers. Our specific model of the hippocampus was based upon early CLS models (K. A. Norman & O’Reilly, 2003; O’Reilly & Rudy, 2001), with additions of theta-phase dynamics (Ketz et al., 2013) and EDL from the entorhinal cortex input layer (ECin) to CA3 (Zheng et al., 2022). The main changes in the present model are the expanded notion of temporal context into various spectra following evidence for this in EC (Bright et al., 2020; Tsao et al., 2018; Umbach et al., 2020) and the continuity of time across different learning epochs and tests. For this reason, we call this the HipSTeR (Hip-pocampus with Spectral Te-mporal Representations) model. Upon publication, please see https://github.com/ccnlab/hipster for all code and detailed explanations, including annotations, fully documented equations, and example simulations, including the model.

Temporal representations were divided into pools that shared a common underlying rate of drift, which involved random bit flips in activation/deactivation at each time step (Estes, 1955a). In this way, time was translated into a spatial code that changed at each moment (Buonomano & Merzenich, 1995). To equalize the overall level of activation over time, whenever an active neuron became inactive, a random, previously inactive neuron became active. Following evidence that other EC cells (grid cells) have discrete spatial frequencies (Stensola et al., 2012; Wei, Prentice, & Balasubramanian, 2015; Whittington et al., 2020) and that drifting EC cells may have discrete drift rates (Aghajan et al., 2022), drift rates spanned a discrete set of values along a spectrum of timescales separated by powers of 2, with 1/4 (2^2^) of the neurons in the fastest pools flipping at each time step and 1/512 (2^9^) in the slowest drifting pools. Differences between the imposed drift rate and actual drift at each time step were carried over into successive time steps so that, especially in the slowest pools with drift rates of less than one unit per time step, drift would eventually occur at an approximately consistent rate across time. Figure 2A depicts the representational drift within each pool via its respective auto-correlation (Pearson *r*) against an initial time point (t = 0).

**Figure 2:**
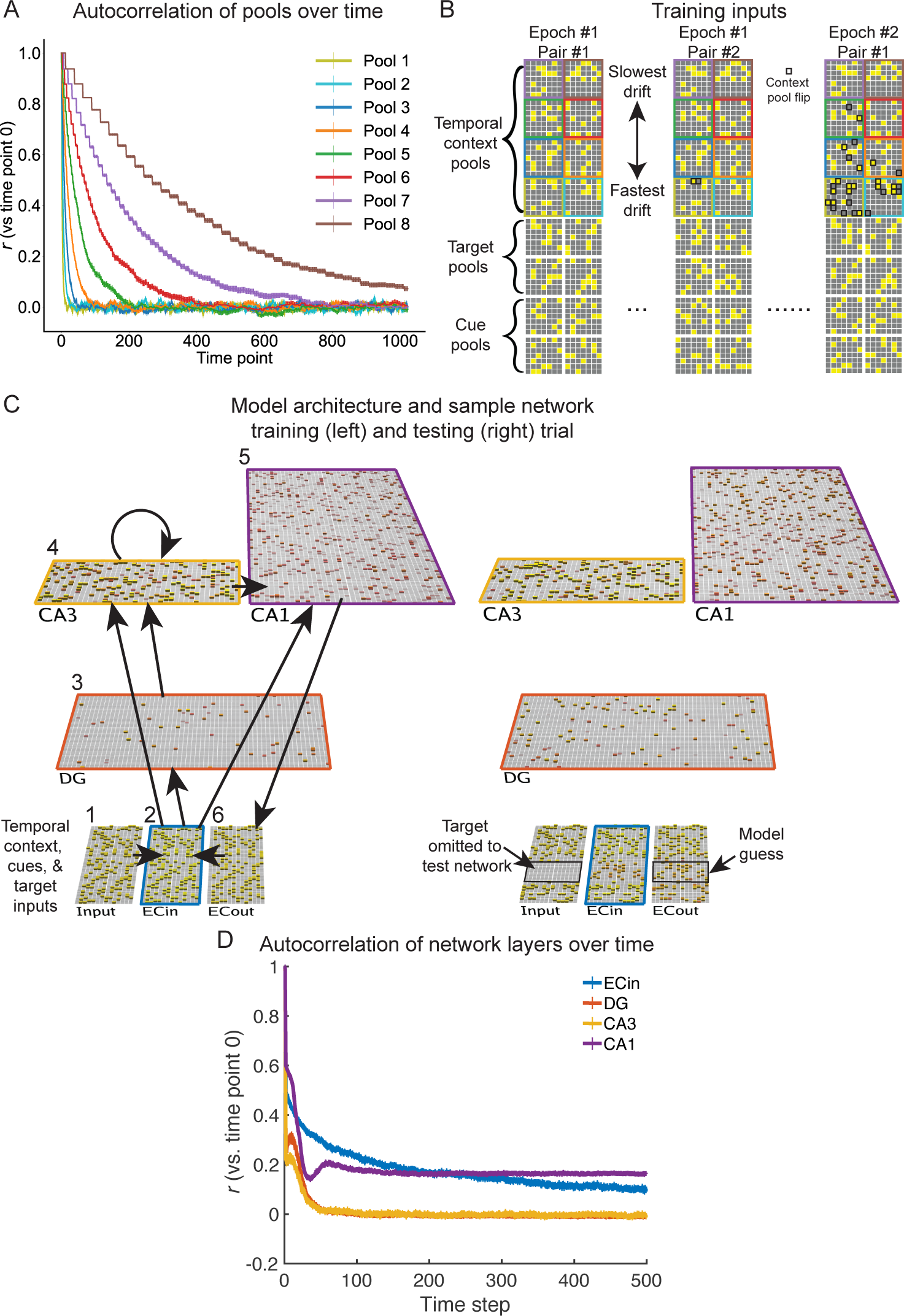
Temporal representations and their implementations in HipSTeR. (A) Temporal representations were separated into different pools of activity that simulated relatively independent cortical inputs and drifted at different rates. Autocorrelations of each temporal context pool were plotted over time against their initial time point. (B) Inputs to our complementary learning systems (CLS) model comprised the eight temporal context pools and four pools each representing cues and targets for paired associates learning. Shown are two successive cue-target pairs during a first training epoch and the first pair again in a second training epoch. Black squares indicate flips in the context pools, showing how drift proceeds differently across pools. (C) Training inputs from (B) entered HipSTeR via ECin, after which they entered the hippocampal loop, with all model connections depicted by arrows on the left. During training (left), the model learned the input patterns. During testing (right), it was presented with the cue and current temporal context pools but without the target pools. Performance was measured by how well the model reproduced target pool activity in ECout. Numbers correspond to layer numbers in the detailed explanations within the Methods-Model Architecture section. (D) Autocorrelations for the primary network layers are depicted as in (A) as the correlation of the activation pattern in each layer against its pattern on the first time step. This was run on a separate, “pure” version of the model without any cue-target repetitions, meaning there were 500 unique cue-target pairs alongside identical temporal context pools. Colors correspond to those surrounding each network layer in (C).

Drift was implemented both within lists across successive trials and between lists (Figure 2B). Within lists, successive trials always occurred after one time step, so the full list of 16 paired associates always spanned 16 time steps. Accounting for the idea that focused tasks or events create some stable state (Antony et al., 2021; Baldassano et al., 2017), after which larger “shifts” in activity occur (DuBrow, Rouhani, Niv, & Norman, 2017), within-list drift was set to 1/4 the rate of normal, between-list drift. Between-list drift occurred to varying extents depending on the experimental condition except in the No Drift comparison condition.

The HipSTeR architecture (Figure 2C) involved the following layers:

1. An input layer comprising 16 pools of 49 neurons each: four cue pools, four target pools, and eight temporal context pools (Figure 2B). These pools were organized separately to reflect how information coming from various cortical regions converges in the hippocampus (Eichenbaum et al., 2007). This layer has only direct, one-to-one forward connections with ECin. Additionally, this arrangement of separating temporal context from other inputs reflects the fact that multidimensional representations of global drift over time versus externally driven contextual factors (e.g., spatial environment) remain largely orthogonal (Keinath & Brandon, 2022).
2. An ECin layer, which receives these signals from the input layer and sends connections into DG, CA3, and CA1 of the hippocampus. ECin, which resembles superficial EC layers (Witter et al., 2017; Zhang et al., 2023), also receives inputs from ECout, which serves as the initial output layer from the hippocampus and resembles deep layers of EC. ECin projections to DG and CA3 via the perforant path are broad and diffuse with a 25% chance of connection.
3. A large DG layer, which features high inhibition (Coulter & Carlson, 2007; O’Reilly & Munakata, 2000), resulting in only very sparse representations that functionally separate the patterns of similar inputs from ECin. In HipSTeR, DG has an inhibitory conductance multiplier of 2.9, resulting in activity of about 1% of neurons. In turn, DG sends outputs to CA3 via strong mossy fiber projections (Henze, Wittner, & Buzsáki, 2002; Vyleta, Borges-Merjane, & Jonas, 2016), which have a multiplier of 4 and give it a stronger influence on CA3 activity than the more direct perforant path inputs from ECin.
4. A CA3 layer, which receives inputs from both ECin and DG and projects to itself (via recurrent collateral connections) and to CA1. The recurrent collaterals – which are fairly strong, with a strength multiplier of 2 in the model – have been theorized to be important for pattern completion because an activated representation can retrieve its previously learned association within this layer (Marr, 1971; O’Reilly & McClelland, 1994).
5. A CA1 layer, which receives and compares input from ECin and CA3, therefore serving as the convergence point for two hippocampal pathways and sends information back out of the hippocampus to the entorhinal cortex output layer (ECout). The pathway from CA3, commonly referred to as the trisynaptic pathway (ECin *→* DG *→* CA3 *→* CA1 *→* ECout), essentially separates common inputs, binds items to contexts, and completes previously stored patterns based on degraded inputs. Evidence for rapid trisynaptic learning comes from findings such as the high inhibition featured in DG, which allows for a separation between highly similar pattern inputs (Leutgeb & Leutgeb, 2007; Vazdarjanova & Guzowski, 2004), the importance of area CA3 in learning new paired associates (Rajji, Chapman, Eichenbaum, & Greene, 2006), and the role of these regions together in discriminating between highly similar information in memory (Bakker, Kirwan, Miller, & Stark, 2008). Note that here we suggest an important role for a disynaptic, ECin *→* CA3 *→* CA1 *→* ECout sub-pathway, following modeling results that this pathway can support generalization (Kang & Toyoizumi, 2023; Kowadlo, Ahmed, & Rawlinson, 2019) and learns via EDL (Zheng et al., 2022). The pathway from ECin *→* CA1 *→* ECout constitutes the monosynaptic pathway of the hippocampus (Schapiro et al., 2017), which allows CA1 to directly encode target ECin activity (Grienberger, Magee, & Duncan, 2022) and sends activity from the hippocampus back into cortex. Evidence for slower monosynaptic learning comes from its having a slightly slower learning rate (Nakashiba, Young, McHugh, Buhl, & Tonegawa, 2008) and more overlapping, generalized representations (Fenton et al., 2008; Leutgeb et al., 2004; Schapiro et al., 2017; Singer, Karlsson, Nathe, Carr, & Frank, 2010). These connections remain within pools, following their point-to-point anatomical connectivity patterns (Witter et al., 2017). This pathway therefore serves an autoencoder function, which translates the pattern-separated representations from the trisynaptic pathway back into a common reference frame for the cortex.
6. An ECout layer, which serves as the output of the hippocampus and therefore the hippocampal network’s “guess” during testing (Figure 2C). Additionally, it also serves as the input back into ECin, which can result in different activations in successive cycles through the hippocampus (Kumaran & McClelland, 2012; Schapiro et al., 2017).

The effects of drift differed across layers of the network. We depicted this without any cue-target pair repetitions by training the network on different cue-target associations for each of 500 time steps and calculated the autocorrelation of each region with its initial time step. Of the hippocampal areas, CA1 showed the slowest pace of drift, followed by DG, which was followed very closely by CA3 (Figure 2D).

Model training and testing followed four discrete phases resembling activity during the four quarters of the hippocampal theta rhythm (Ketz et al., 2013). The model learned via two EDL mechanisms. In the first mechanism, the first three quarters constituted what are considered the minus phases, whereby the network produced an expected output based on its weights and input activations. The fourth and final quarter was the plus phase, whereby the target activation was provided from ECin *→* ECout and thereby learning occurred based on the difference between the network’s prediction from the minus phases into ECout and the actual outcome. The first and fourth theta phases came during theta troughs, when CA1 was strongly influenced by ECin (Siegle & Wilson, 2014). Conversely, at theta peaks, CA1 was strongly influenced by CA3, which involved a guess based on activations and previously stored patterns. During the plus phase, ECin drove both CA1 and ECout activity, effectively clamping the correct answer in both EC layers and forcing weight adjustments in CA1. Therefore, across learning, ECout activity came to resemble ECin activity via the CA1 projection during the minus phases (without the direct ECin *→* ECout input). The second mechanism involved EDL in CA3 (Zheng et al., 2022). This error arose as a form of temporal difference learning between different pathways converging on CA3 neurons (Sutton & Barto, 1998): direct input from ECin (via the perforant path) and CA3 recurrent collateral activations arrived on CA3 neurons within the first quarter of the theta cycle, and critically, this input preceded signals from the multisynaptic ECin *→* DG *→* CA3 pathway (Yeckel & Berger, 1990). This minus phase constituted CA3 activity prior to DG inputs and the plus phase occurred when they arrived. Therefore, the pattern-separated DG activation acted as a teaching signal to correct the predicted pattern in CA3 based on perforant path + recurrent collateral activations alone (Kowadlo et al., 2019).

In our simulations, as in prior models (Ketz et al., 2013; Zheng et al., 2022), temporal context drift occurred within trials of an epoch. However, drift differed in HipSTeR in multiple ways. As mentioned above, drift occurred in all simulations across a spectra of time constants in a manner that was constant within each temporal context pool. Additionally, we differentially modified drift in some experimental conditions in two other ways. First, drift often occurred between learning epochs, with the number of drifting time steps defined as those coming after the final learning trial of an epoch and before the first trial of the next epoch. We refer to this drift as the ISI. The exception to this was the No Drift condition, in which drift still occurred within-epoch, but each epoch of training was identical. Given that neurobiological drift occurs [e.g., Tsao et al. (2018)], such a condition is biologically impossible, as an agent would never return to the exact same drifting neural pattern during re-learning. Note that similar neural firing patterns can recur even on long timescales with repeated experiences (Liu et al., 2022; Sun, Yang, Martin, & Tonegawa, 2020), but they are not identical. However, it is common to train neural networks this way, including in prior versions of the CLS architecture, so we used the No Drift condition as an illustrative comparison.

Second, drift often occurred after the final training epoch and before the model was tested. We refer to this as the RI, following the labeling convention used in human behavioral experiments employing memory tests after various intervals. We defined the number of drifting time steps as those occurring after the final learning trial of the final training epoch and before the first trial of testing. Drift continued at the same rate for each trial of the testing epoch. Similar to the No Drift training condition, we also had a No Lag (RI-0) retention interval, for which the temporal context given at test was the exact temporal context used in the final training epoch. Similar to the No Drift condition above, the No Lag condition is biologically impossible, but likewise served as a useful control to assess the network’s ability to recall under the exact conditions of at least one of its learning epochs.

HipSTeR has trisynaptic (ECin *→* DG *→* CA3 *→* CA1 *→* ECout), disynaptic (ECin *→* CA3 *→* CA1 *→* ECout) and monosynaptic (ECin *→* CA1 *→* ECout) pathways. Thus, it is possible to turn off learning (prevent all weight changes) in some pathways and reasonably expect some alternate learning to proceed (Schapiro et al., 2017). Additionally, ECin *→* CA3 learns via both Hebbian learning and EDL, which changes weights proportionally to the difference between CA3 activity with ECin, DG, and recurrent CA3 inputs present against activity prior to DG input. We can therefore isolate the importance of these learning rules within ECin *→* CA3 by turning off EDL while leaving Hebbian learning intact or turning off all learning. Altogether, we compared the full HipSTeR model to alternative models in which learning pathways were affected in the following ways: ECin *→* DG (no learning), ECin *→* CA3 (no learning), ECin *→* CA3 (no EDL, but Hebbian learning present), ECin *→* CA1 (no learning), CA3 *→* CA3 (no learning), and CA3 *→* CA1 (no learning). To assess the impact of multi-scale drift versus other uniform drift implementations, we also compared simulations of our control HipSTeR model against networks wherein all temporal context pools drifted at a uniform slow, medium, or fast rate. The slow-drifting network used the slowest drift rate from HipSTeR (1/512 per time step), the fast-drifting network used the fastest drift rate (1/4), and the medium-drifting network used a medium rate (1/64).

### Experimental conditions in spacing effect simulations

We will now outline specific simulations of prior behavioral findings. For these and other simulations, our hypotheses were not pre-registered. More information about the original studies can be found in the corresponding region in the Results section. To model spacing effects with one variable ISI (Cepeda et al., 2008), we used four training epochs with a unique scheduling procedure. The first two epochs simulated the encoding and one perfect recall trial of the first learning session. These were implemented in direct temporal succession (ISI = 2 between lists) in the model. The third and fourth epochs simulated the two practice + feedback trials of the second learning session, and these also occurred in direct temporal succession (ISI = 2). Critically, the ISI between the second and third lists differed across experiments. These 8 ISIs spanned powers of 2, from 4 (2^2^) to 512 (2^9^). We then used four RIs after the final training epoch spanning powers of 2, from 64 (2^6^) to 512 (2^9^). We chose these ISI and RI combinations so that we had a mix of ISI:RI ratios, ranging from far less to far greater than 1. We fit these data to find optimal ISIs using an equation from (Cepeda et al., 2008): *y* = *−a ∗* (*log*(*x* + 1) *− b*)^2^ + *c*, where *y* is recall performance, *x* is ISI, and *a*, *b*, and *c* are free parameters. From the best-fit parameters, we found the time point corresponding to maximum performance in the curve to obtain the optimal ISI.

For the spacing override effect (Rawson, Vaughn, Walsh, & Dunlosky, 2018), three experimental groups had three epochs of training with lags of 2 (Low spacing), 8 (Medium spacing) or 32 (High spacing). Following this initial training session were two training epochs separated by 128 time steps each. After these epochs, the final test occurred after another 128 time steps. Note that the Low and High spacing conditions approximately map onto the Rawson et al. (2018) conditions of Lag-15 and Lag-47 because they used different naming conventions; by their conventions, our Massed would be Lag-17 (15 drifting time steps during the training list itself + 2 time steps after the list) and our Lag-32 would be Lag-47 (15 during training + 32 after the list).

To model the importance of absolute amounts of spacing and different spacing regimens, including expanding, contracting, and equal spacing (Küpper-Tetzel et al., 2014), we used four different experiments, each using five training epochs with unique ISIs. All experiments had very short intervals between the first, second, and third training epochs. Following this, the first three experiments had intervals between training epochs that were Expanding (16, 256), Contracting (256, 16), or equally spaced intervals (136, 136) that matched the overall drift of the prior two (Equal, match). To demonstrate the importance of absolute spacing, the final experiment (equal, compressed) used equally spaced intervals between training epochs but 1/8 of the amount of overall spacing (17, 17). After the final training epoch, retention intervals occurred after drift corresponding to values spanning powers of 2, from from 32 (2^5^) to 2048 (2^11^) time steps.

To model list-wise spacing with repeated variable ISIs (Bahrick et al., 1993), we used five training epochs using consistent ISIs in each experimental condition. The drifting conditions had ISIs of powers of 2, including 8 (2^3^), 64 (2^6^), and 512 time steps (2^9^). After the final training epoch, RIs occurred after drift corresponding to values of powers of 2, including 64 (2^6^) and 1024 (2^10^). Additionally, a Scrambled condition used scrambled temporal context vector pools. These pools were only scrambled once after training, meaning that they used the same drift rate during the testing epoch, which controlled for the presence of drift during testing.

In later simulations aimed at demonstrating the mechanisms of learning in HipSTeR, we primarily used the approach from modeling Bahrick et al. (1993) (repeated variable ISI) with a few additions. First, we added more ISIs and RIs. ISIs spanned powers of 2, from 2 to 512 (2^9^) time steps, while RIs went from 32 (2^5^) to 2048 (2^11^) time steps. Second, we added a No Drift ISI condition that involved training the model using the same temporal context vectors in each epoch. Drift still occurred within this list, but the input pattern for e.g., the first cue-target pair was the exact same across epochs. Third, we added a Scrambled ISI condition, whereby we completely scrambled the temporal contexts before each new training epoch while preserving within-list drift within a training epoch. Finally, we added a No Lag RI condition that was tested using the temporal context vector from the final training epoch. We still used the Scrambled RI condition as in the Bahrick et al. (1993) simulations that involved scrambling the temporal contexts before the final test.

### Measuring model memory performance, CA3 error, representational similarity, and weights

In each experimental condition, model weights were re-initialized and trained anew with random weights within what we call runs. Different runs were analogous to random assignment for human participants, and we therefore performed inferential statistics across different runs of the model.

We tested model memory performance by assessing the activation of ECout neurons after the minus phases against the plus phase. Neuronal activity above and below 0.5 was considered active and inactive, respectively. For correct performance on a given trial, the proportion of active neurons that were expected to be inactive and the proportion of inactive neurons that were expected to be active had to both be below 0.2.

To assess error across training epochs, we calculated the absolute difference in CA3 activation in the first quarter (Q1) versus the final plus phase (Q4) and divided this quantity by the average trial-wise CA3 activation. This error metric indicated how easily the network produced the intended output. Given that it generally scales with poor current performance but greater subsequent learning, it provides a useful proxy for the concept of desirable difficulties in learning, in which making learning more difficult often has positive long-term consequences for memory retention (R. Bjork & Bjork, 1992).

To gain a sense of how representations changed across epochs, we calculated the representational similarity in CA3 during all training trials. To do this, we correlated across-CA3 neuron activation patterns at the end of each trial in the current epoch (starting in epoch 2) against each trial in the prior epoch. These values were then separated based on whether they involved the same versus different input cues.

To assess the structural changes to the network across learning and as a function of experimental condition, we measured the average weight strength between ECin *→* CA3 neurons for each pool. We measured changes in this pathway (the perforant path) because of its role in supporting cue-target learning and its EDL properties. To assess how various experimental conditions affected representations in different temporal pools, we calculated the average strength from neurons in each ECin temporal pool separately. To assess the effects of training regimens, we used values after the final training epoch and contrasted various experimental conditions (e.g., Drift vs. No Drift) across runs.

### Experimental conditions in item-wise and block-wise decontextualization simulations

Following evidence of decontextualization in the spacing effect simulations, we simulated more canonical decontextualization paradigms using epoch-wise (Imundo et al., 2021; S. M. Smith et al., 1978) or item-wise contexts (S. M. Smith & Handy, 2014, 2016). Rather than 8 temporal context pools, our inputs here involved 6 temporal context pools and 2 other context pools representing either epoch-wise or itemwise context. For epoch-wise contexts, which were analogous to learning associations for an entire session within a spatial context (Imundo et al., 2021; S. M. Smith et al., 1978), the other context pools were either the same for all training epochs (Constant Epoch condition) or new for each epoch (Variable Epoch condition). For trial-wise contexts, which were analogous to having some incidental background context present during each individual learning association (S. M. Smith & Handy, 2014, 2016), the other context pools were unique for each cue-target pair but were either the same across each training instance of that pair (Constant Trial) or changed each time (Variable Trial). Finally, the other context pools at test could either be the same as the context from the first training epoch (Old Test) or a new context (Novel Test). All context vectors were randomly generated and bore no resemblance to others. For these experiments, we used a short ISI (2 steps) and moderate RI (512 steps) for all simulations to control for the temporal context.

## Results

We began our investigations by simulating a number of behavioral findings from the spacing effect literature. Following this, we probed the mechanisms by which our HipSTeR model learned amid constantly drifting inputs during training, including assessments of error, representational similarity within layers of the network, weight changes, eliminating learning in specific HC pathways under various training regimens, and comparing our multi-scale drift model to alternative models with uniform drift. Lastly, to connect our findings to more canonical ideas of decontextualization, we simulated decontextualization paradigms that were unrelated to temporal context.

### Optimal ISI decreases with RI and optimal ISI:RI ratio decreases with increasing RI: list-wise spacing with one variable ISI

We first simulated an experiment (Cepeda et al., 2008) containing a single variable ISI separating two training sessions. Briefly, Cepeda et al. (2008) had human subjects learn lists of paired associates to a criterion of two correct trials, wait a variable number of days before performing to a criterion of two correct trials [0 (3 min), 1, 2, 7, 21, or 105 days], and testing thereafter after RIs of 7, 35, 70, or 350 days. They found that the relationship between ISI and memory recall was non-monotonic (e.g., optimal at some ISI between the extremes of those tested) and that the optimal ISI increases with RI (Figure 3A). Additionally, they found an intriguing relationship between ISI and RI, in that the optimal ISI:RI ratio is not consistent but actually decreases with increasing ISI.

**Figure 3:**
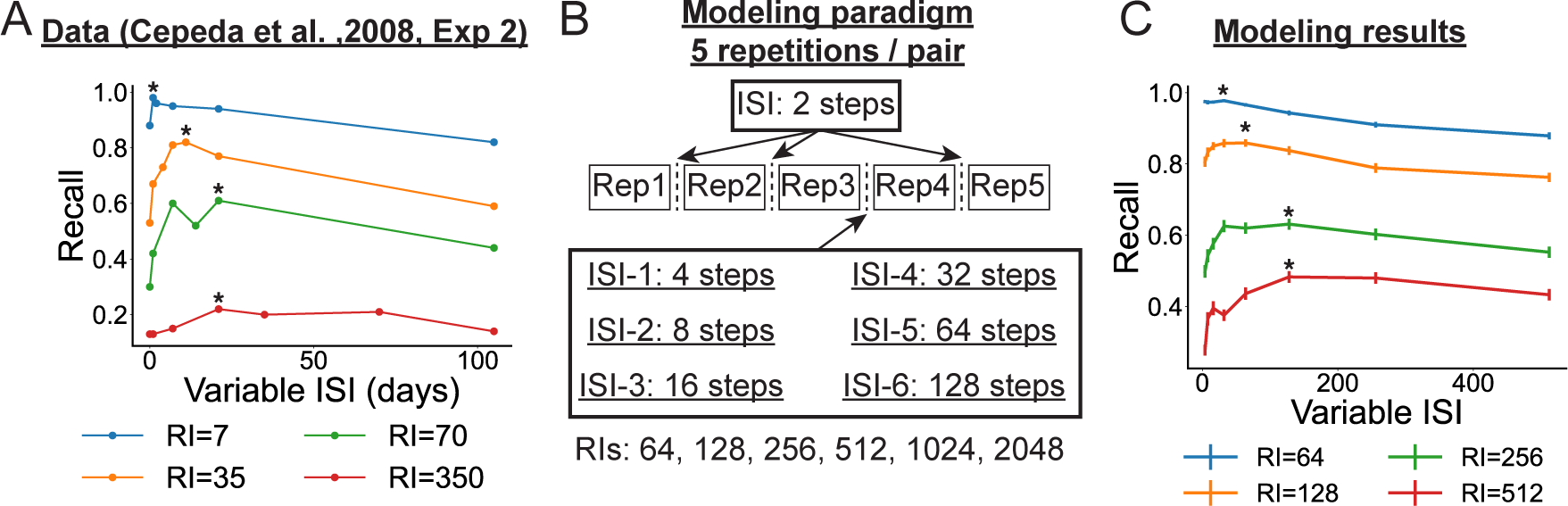
Simulations showing optimal ISI depends on RI and optimal ISI:RI ratio decreases over time. (A) Data replotted from Cepeda et al. (2008). (B) Our modeling paradigm, wherein only a single ISI between repetition 3 and 4 differed across the conditions. Note that steps are used as a proxy for time in the behavioral experiments. (C) Modeling results. Data from simulations were plotted as mean ± SEM across runs in the model. In (A) and (C), the optimal ISI for the same RI is marked with an asterisk.

We simulated these results by (1) training the model to learn lists in short succession (to capture initial study and one round of learning to criterion), (2) imposing one variable ISI [from 4 (2^2^) to 128 (2^7^) trials] between the second and third training epochs, (3) training the model on two more lists occurring within short succession (to capture the final rounds of learning), and (4) testing the model after variable RIs [from 64 (2^6^) to 512 (2^9^) steps] (Figure 3B). First, we performed a 2-way, ISI x RI between-subjects ANOVA, and we found a main effect of RI (*F*(3,3564) = 2962.0, *p <* 0.001), a main effect of ISI (*F*(8,3564) = 20.5, *p* = 0.19), and a significant interaction (*F*(24,3564) = 67.1, *p <* 0.001). Second, similar to Cepeda et al. (2008), we found that the relationship between ISI and RI was non-monotonic, peaking at medium (not the shortest nor the longest) ISIs for each RI. Third, the optimal ISI increased with increasing RIs; to find this, we used model fits based on a three-parameter equation from Cepeda et al. (2008), *y* = *−a∗*(*log*(*x*+1)*−b*)^2^ +*c* (see Methods for details). The optimal ISIs from these model fits were 14.3, 34.8, 70.7, and 204.1, respectively. Third, the ISI:RI ratio with increasing RI decreased, as these ratios were 64/14.3 = 4.48, 128/34.8 = 3.68, 256/70.7 = 3.62, and 512/204.1 = 2.5, respectively (Figure 3C). Therefore, we captured the main principles of the spacing effect outlined in Cepeda et al. (2008). Of these principles, the first is especially important conceptually because it suggests there is not one solo factor underlying memory strength – if there were, the ISI condition resulting in the greatest strengthening would produce the best memory performance regardless of RI. We will later demonstrate that these differentiating factors depend on the amount of overlap between the temporal context at test and the learned contexts (as governed by the RI), the strength of each of the weights in each of the layers, and the direct strength between the cue and target pools.

### Re-learning override occurs with relatively large amounts of spacing

If a memory has been learned with low spacing, can it still benefit from spacing later? Next, we addressed this question, following what has previously been referred to as the re-learning override effect (Rawson & Dunlosky, 2011; Rawson et al., 2018). Briefly, this effect occurs when an initial, small difference in either spacing (Rawson et al., 2018) or initial learning (Rawson & Dunlosky, 2011; Rawson et al., 2018) becomes largely (but not necessarily completely) overridden by re-learning after a longer spacing interval. That is, the relative gain after a longer spacing interval is larger for an initially weaker memory, whether it be weaker because of fewer initial learning trials or more massed training. To demonstrate the override effect in terms of initial differences in spacing, we turned to the learning criterion = 3 conditions of Experiment 1 in Rawson et al. (2018). In this condition and experiment, subjects initially studied Lithuanian–English word pairs with different initial lags of either 15 or 47 during the first session. They were then practiced in the same order until they were retrieved correctly thrice. After this session, subjects returned for a series of re-learning sessions spaced one week apart (Figure 4A, right). The relative gain when re-learning after a large temporal gap was larger for initially less-spaced memories.

**Figure 4:**
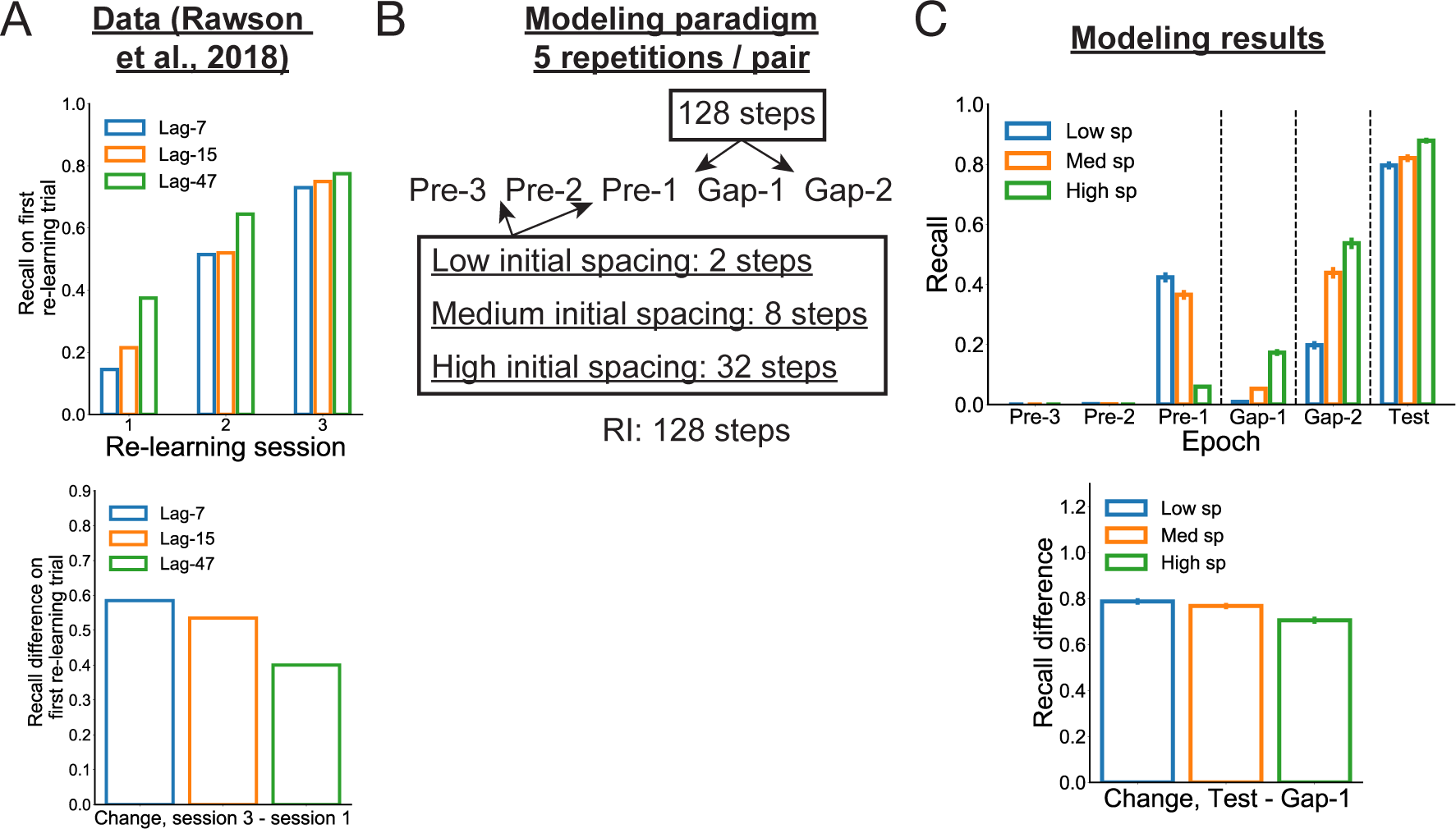
Simulations showing that re-learning override occurs when later spacing is greater than initial spacing. (A) (top) Data replotted from Rawson et al. (2018). (bottom) Recall differences from first to third re-learning sessions showed larger benefits in initially lower spacing conditions. (B) Our modeling paradigm, wherein we created Low, Medium, and High initial spacing conditions before longer gaps before re-learning epochs and a final test. (C) Modeling results. (top) Data from simulations were plotted as mean ± SEM across runs in the model, including performance on initial training runs (left), after larger gaps (middle), and at final test (right) (in some cases, error bars are too small to visualize). (bottom) Model recall differences from just after the first large gap (Gap-1) to the final test showed larger benefits with initially lower spacing.

To model the spacing override effect, we created three experimental conditions (Figure 4B). These conditions used three initial epochs of training with lags of 17 (15 for training list + 2 extra time steps) (Low spacing), 23 (15 for training list + 8 extra time steps) (Medium spacing), and 47 (15 for training list + 32 extra time steps] (High spacing). In all conditions, initial training was followed by two later training epochs, each after 128 time steps, followed by a final test after another 128 time steps.

Investigating memory at each training epoch in the first two groups, we first found a typical effect of faster learning in the lower spacing conditions before the first larger gap (final training epoch, *F*(2,297) = 194.2, *p <* 0.001) (Figure 4C). After this gap, memory was superior for the higher spacing conditions (*F*(2,297) = 122.6, *p <* 0.001), in line with classic spacing effects. However, after two large gaps, memory benefits were higher for the initially low spacing groups at the final test (*F*(2,297) = 9.0, *p <* 0.001; follow-up *t*-tests for the low versus and medium and high spacing groups indicated both *t >* 3, *p <* 0.003), demonstrating a re-learning override of initial, relatively small spacing.

In some ways, the override effect of initial learning criteria is surprising, as it has long been known that increasing the number of learning trials slows the rate of forgetting [e.g., Krueger (1929)]. However, these benefits tend to be weak and transitory when overlearning occurs within close temporal succession (or all within the same session) (Driskell, Willis, & Copper, 1992; Elliott, Isaac, & Muhlert, 2014; Pyc & Rawson, 2009). Therefore, the relative change in memory change offered by later spacing is substantial enough to drastically reduce or eliminate these initial differences. These findings underscore the idea that learning that is relatively compressed in time *overfits* to one temporal context that ultimately proves unhelpful when that temporal context is no longer active. It also points to the importance for long-term memory of absolute spacing, or the total time between the first and last training instances for a given memory, which we will cover in more detail in the following section.

### Alternative schedules and absolute spacing: contracting, expanding, and equally spaced intervals

The re-learning override effects of large spacing suggest that the greatest temporal determinant of later memory may be the absolute amount of spacing between all training instances (Karpicke & Bauernschmidt, 2011). However, a number of investigations have examined the importance of alternative training schedules, such as ISIs that expand, contract, or remain equal across training instances (Gerbier, Toppino, & Koenig, 2015; Mettler, Massey, & Kellman, 2016; Toppino & Gerbier, 2014; Toppino, Phelan, & Gerbier, 2018). In one study (Küpper-Tetzel et al., 2014), subjects learned paired associates before re-learning on multiple sessions after either expanding (1-, 5-, and 9-day), contracting (9-, 5-, and 1-day), or equal (all 5-day) ISIs. Final tests were given after either 1, 7, or 35 days. Free recall performance (requiring both items of a pair to be recalled and matched) showed that the contracting schedule was superior to the equal and expanding schedules for the early (1 and 7 day) RIs, whereas equal and expanding schedules were superior to the contracting schedule at long RIs (Figure 5A). (They focused on free recall because cued recall was at or near ceiling performance, though we will model cued recall.)

**Figure 5:**
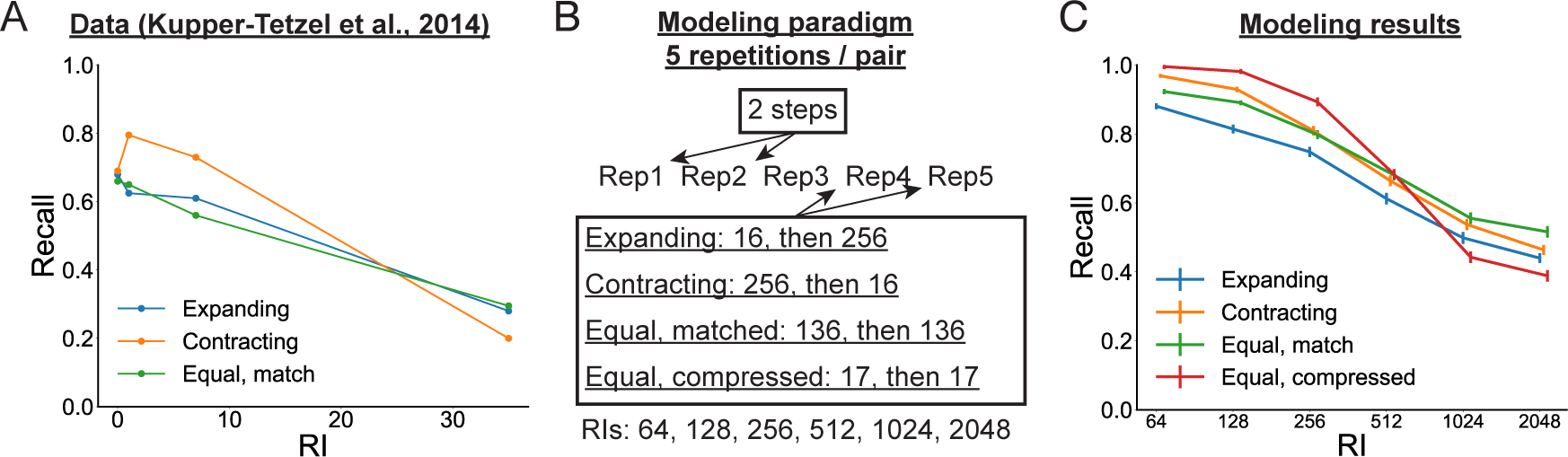
Simulations showing the importance of absolute spacing and investigations of alternative schedules. (A) Data replotted from Küpper-Tetzel et al. (2014). (B) Our modeling paradigm, wherein we created Expanding, Contracting, Equal and Matched, or Equal and Compressed conditions to show the impact of different re-learning schedules and absolute spacing. (C) Modeling results. Data from simulations were plotted as mean ± SEM across runs in the model (in some cases, error bars are too small to visualize).

We modeled the effects of alternative schedules and absolute spacing using four experimental conditions with unique ISIs (Figure 5B). Here the intervals before the two final training epochs differed in the following conditions by the number of drifting time steps: Expanding (16, 256), Contracting (256, 16), Equal matched (136, 136) and Equal compressed (17, 17). The first three conditions allowed us to assess the importance of alternative schedules, while the final condition allowed us to again assess the importance of absolute spacing. Finally, RIs occurred after 32 (2^5^) to 2048 (2^11^) time steps.

Overall, a two-way, learning schedule (Expanding, Contracting, Equal, matched, or Equal, compressed) x RI (32 to 2048) ANOVA revealed significant main effects of schedule (*F*(3,2376) = 21.7, *p <* 0.001), RI (*F*(5,2376) = 671.2, *p <* 0.001), and an interaction (*F*(15,2376) = 16.8, *p <* 0.001). Similar to Küpper-Tetzel et al. (2014), we found that the Contracting schedule was superior performance to the other conditions at the early retention interval: the order, considering significant differences at *p <* 0.05 as *>*s and insignificant ones as =s, was Equal, compressed *>* Contracting *>* Equal, matched *>* Expanding (Figure 5C). We believe that this occurred because this group had the most training opportunities within a short temporal interval of these tests, allowing for better training within this narrow temporal context. However, at the longest RIs, we found that this advantage had disappeared, reversing against the Equal condition and showing no difference from the Expanding condition: the order was Equal, matched *>* Contracting = Expanding (*p* = 0.87) *>* Equal, compressed. Note that the insignificant differences between Expanding and Contracting at long RIs differs from the Küpper-Tetzel et al. (2014) results for their longest RI. All three groups showed superior memory at the longest RIs against the Equal Compressed group (all *p <* 0.023), once again highlighting the critical importance of absolute spacing when tests occur after long RIs.

Therefore, we showed that contracting schedules had superior performance at short RIs, likely due to their having more training examples within close temporal proximity of these tests, but this advantage disappeared over time. Additionally, the Equal spacing condition produced superior recall at some long RIs. One aspect of the Küpper-Tetzel et al. (2014) results we were not able to capture was superior memory for expanding than contracting schedules at the longest time points. Benefits for expanding over contracting schedules are not always found at the longest RIs [e.g., Cull (2000); Karpicke and Bauernschmidt (2011)]. However, we believe there is a larger point: the subtle differences in recall across RIs between these three conditions — as well as findings from the re-learning override effects — points to the *absolute* spacing of training instances as the greatest determinant of long-term memory performance. This accords with behavioral findings showing that differences between expanding, equal, and contracting schedules were inconsistent and minor in comparison to differences at three levels of absolute spacing (Karpicke & Bauernschmidt, 2011).

### Spacing effects at extremely long RIs using repeated variable ISIs

Another seminal spacing effect finding involved multiple learning episodes spread over ISIs of up to almost two months and final RIs of up to five years (Bahrick et al., 1993). In the study, four members of the same (Bahrick) family learned foreign language-English pairs over 13 or 26 sessions spaced by 14, 28, or 56 days before a final test one, two, three, or five years after the final learning session. Similar to prior effects showing that short ISIs improve memory at short RIs, performance was best at the end of training for the 14-day interval schedule. However, when assessed after RIs from 1-5 years, performance was best for words in the 56-, then 28-, then 14-day interval (Figure 6A). For our purposes, these findings show that spacing effects can compound over numerous intervals and can still be demonstrated at extremely long RIs.

**Figure 6:**
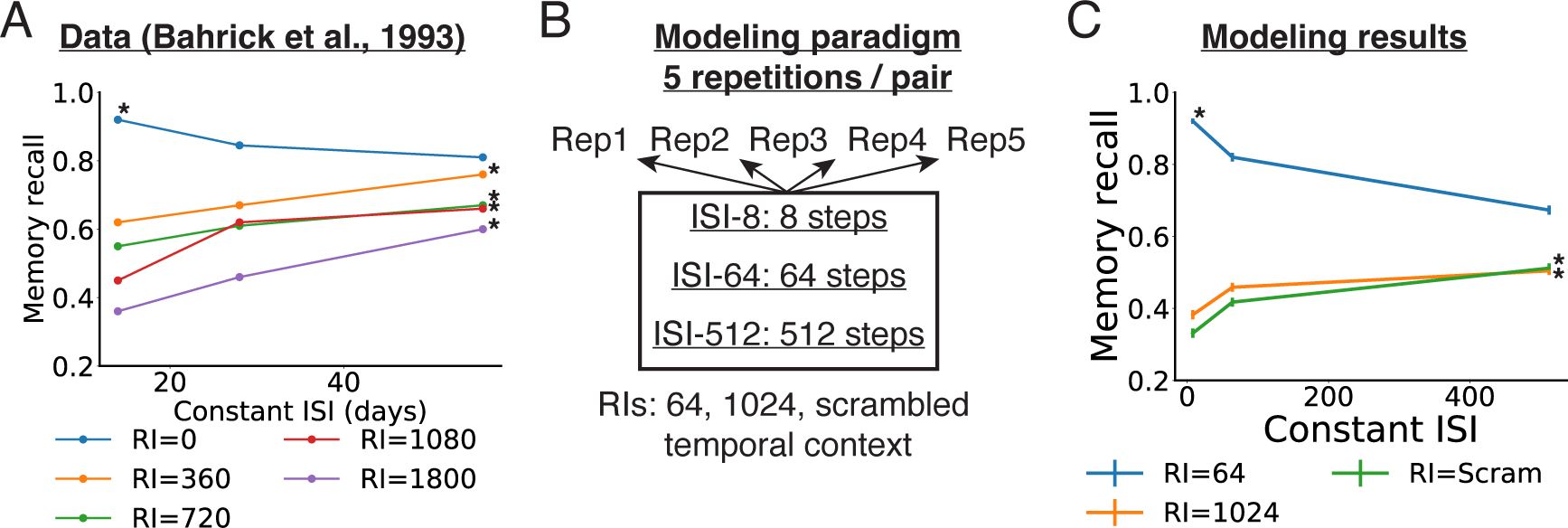
Simulations showing spacing effects at extremely long RIs. (A) Data replotted from (Bahrick et al., 1993). (B) Our modeling paradigm, wherein the same ISI was used between all training epochs and RIs were included at a short and long delay as well as after scrambling the temporal context vector. (C) Modeling results. In (A) and (C), the optimal ISI for the same RI is marked with an asterisk. Data from simulations were plotted as mean ± SEM across runs in the model (in some cases, error bars are too small to visualize).

We modeled these findings by training HipSTeR using the same ISI across each of five epochs, of 8, 64, and 512 time steps (Figure 6B). After the final training epoch, we implemented RIs of 64 and 1024 time steps. We additionally added a condition using scrambled temporal context vector pools to simulate what could arguably occur to temporal context at very long RIs [e.g. 5 years in Bahrick et al. (1993)].

A two-way, ISI (8, 64, or 512 time steps) x RI (64, 1024, or scrambled) ANOVA revealed significant main effects of ISI (*F*(3,888) = 137.7, *p <* 0.001), RI (*F*(2,888) = 1233.5, *p <* 0.001), and an interaction (*F*(6,888) = 109.3, *p <* 0.001). Similar to Bahrick et al. (1993), we found that memory recall in the model was best for the ISI-8, then ISI-64, then ISI-512 at the early RI (Figure 6C) (both *p <* 0.001). For the RI-2048 conditions, model recall followed a ISI-512 *>* ISI-64 *>* ISI-8 (all *p <* 0.001), and for the Scram RI conditions, model recall followed a ISI-512 = ISI-64 (*p* = 0.95) *>* ISI-8 pattern (other contrasts, *p <* 0.001). These results - especially those with a scrambled temporal context - suggest that spacing benefits in HipSTeR go beyond strengthening connections within the temporal context layer, but also benefit *decontextualized* connections between the cues and targets themselves. As we will discuss below in investigating the mechanisms of learning in the model, these benefits accrue during later training epochs based on greater errors between stored and current temporal context vectors.

### Drift vs. no drift: learning mechanisms

Having established that the model can reproduce a number of spacing-related effects on memory, we next probed the mechanisms underlying these effects in HipSTeR. To simplify our initial investigations, we first contrasted performance in the model with no drift between training epochs - the canonical way in which neural networks are trained - against performance with modest drift, specifically four time steps + the drift during the list itself (ISI-4). We tested the model with various RIs after the final training epoch, including a No Lag RI and a condition with a scrambled temporal context (Scram RI) (Figure 7A). As expected, a two-way, ISI x RI ANOVA revealed a significant main effect of RI (*F*(8,1782) = 2135.5, *p <* 0.001), in line with forgetting (Figure 7B). Importantly, the Drift condition outperformed the No Drift condition in recall generally (*F*(1,1782) = 1356.6, *p <* 0.001) and did so increasingly at longer RIs, as shown by an interaction (*F*(8,1782) = 42.2, *p <* 0.001). These results suggest that drift during training makes the learned representations more resistant to further drift. Curiously, and unexpectedly, the Drift condition even slightly bested the No Drift condition at the No lag RI, which featured the exact same temporal context vectors used for all five training epochs in the No Drift condition [*t*(198) = 7.1, *p <* 0.001]. Although these results were unexpected, they are reminiscent of the impoverished learning that occurs in human behavior when two (massed) learning trials are presented with no delay (Benjamin & Tullis, 2010; Thios & D’Agostino, 1976; Xue et al., 2011).

**Figure 7:**
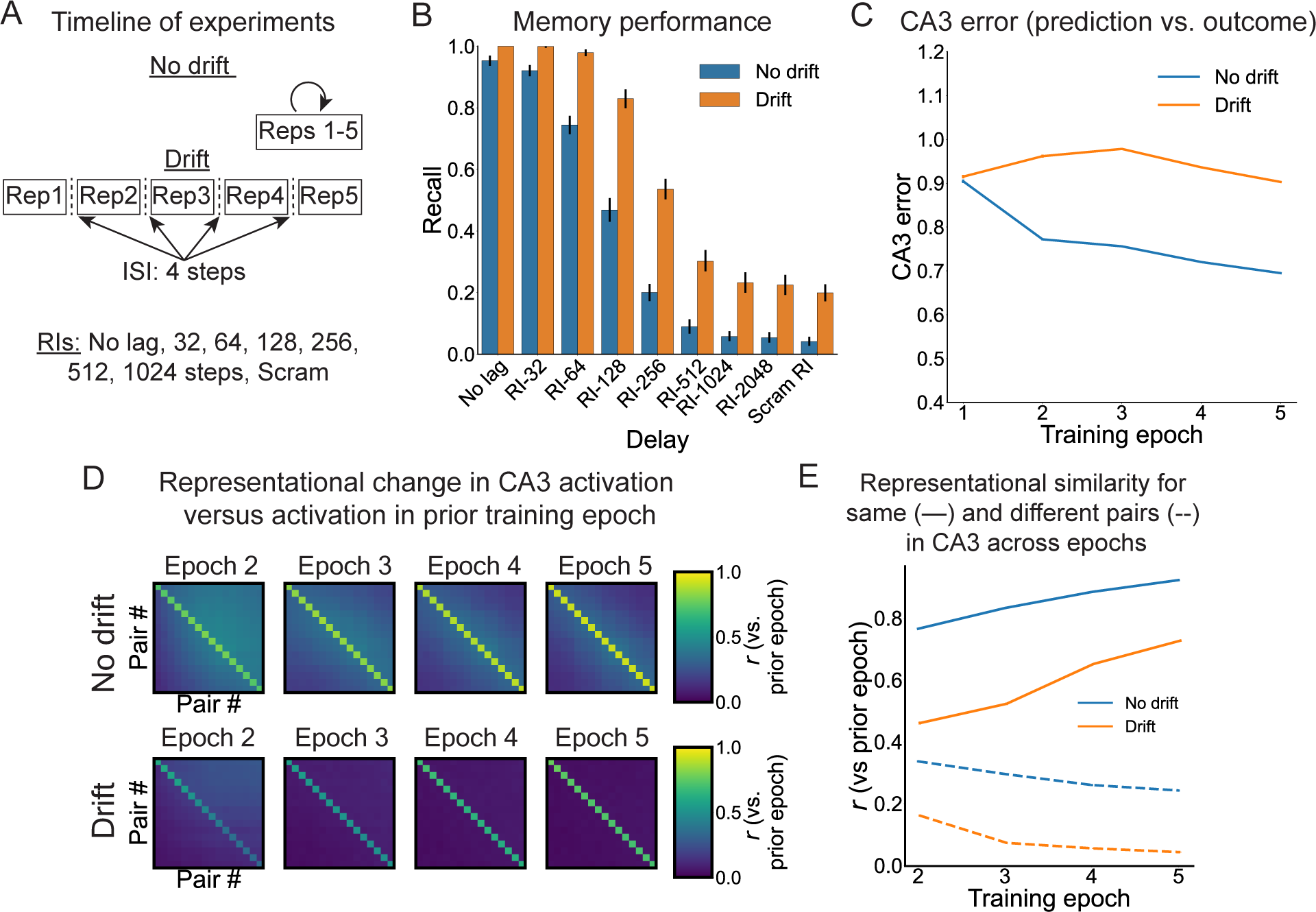
Effects of imposing neural drift between model training epochs. (A) Timeline of experiments including one regimen with five training epochs composed of identical temporal contexts for each paired associate trial (No drift) and another with modest drift between each epoch (4 time points). Each regimen occurred in eight training conditions, followed by retention intervals consisting of either no lag (e.g., the same temporal context as the final training trial) or six lags (32 to 1024 time points). (B) Model memory recall was plotted for No drift and Drift conditions across all retention intervals. Drift improved memory performance, especially after long retention intervals. (C) Error between the first quarter of the theta cycle in CA3 (the model’s prediction based on the stored pattern) versus the final quarter (the “plus” phase) remained higher across training for the Drift than the No drift condition. (D) Representational stability across training epochs, measured by correlating CA3 unit activation patterns for each trial from a given training epoch against each trial of the prior epoch, was plotted for both the No drift (top) and Drift (bottom) conditions. Trials along the diagonal represent the same pairs across epochs, whereas off-diagonal trials represent different pairs. (E) Across training, representations from epoch-to-epoch increased for the same pairs (solid lines) and decreased across pairs (dotted lines). The No drift regime produced higher same- and across-pair representations than the Drift condition. Data in B, C, and E are shown as mean ± SEM across runs in the model (in some cases, error bars are too small to visualize).

We next sought to characterize mechanistic differences between HipSTeR trained in the Drift and No Drift conditions. First, to gain insight as to how EDL differs between these conditions, we measured error in area CA3, which acts as a convergence zone between the direct pathway from ECin and indirect pathway from DG. Recall that training trials in HipSTeR have four quarter stages resembling distinct theta phases. Error in area CA3 occurs because information arrives at different phases from different sources: during the first quarter (Q1), information arrives from ECin via the ECin *→* CA3 pathway, whereas in later quarters, information arrives from ECin *→* DG *→* CA3. As a result, the signal from DG effectively teaches and adjusts direct weights from EC *→* CA3 accordingly. Therefore, we were especially interested in the error contrasting CA3 Q1 and Q4 (plus phase) activity, which drives EDL in this area. In line with typical learning effects when using repetitions of identical input patterns, this error signal quickly abated across training epochs in the No Drift condition (Figure 7C). Conversely, because temporal context patterns continually changed across training epochs in the Drift condition, this error remained high, driving greater subsequent weight changes. These patterns were supported by results from a mixed, two-way ANOVA on CA3 error, with condition (Drift vs. No drift) and learning epoch (1-5) as factors. This ANOVA revealed a main effect of condition [*F*(1,9990) = 5042.5, *p <* 0.001], a main effect of epoch [*F*(4,9990) = 952.1, *p <* 0.001], and a significant interaction [*F*(4,9990) = 623.6, *p <* 0.001].

Next, we asked a related question of how drift affects learning-related representational change on successive training epochs to ask how well learning drives pattern separation across pairs. To do this, we measured activity patterns across CA3 neurons at the end of the trial (Q4), and we correlated each pattern against all pairs in a given training epoch against the prior epoch (e.g., Epoch 2 vs. Epoch 1). To test this, we ran a 3-way, condition (Drift vs. No drift) x learning epoch (2-5) x pattern status (same vs. different) ANOVA. As expected, activity patterns were more similar for same than different pairs in both the No Drift and Drift conditions, as shown by the bright diagonal line in Figure 7D and revealed by a main effect of pattern status (*F*(1,1584) = 153796.0, *p <* 0.001). However, the nature of these correlations differed markedly between the conditions, as revealed by a main effect of condition (*F*(1,1584) = 28344.3, *p <* 0.001). [The main effect of learning epoch, as well as every two-way interaction, and the three-way interaction were all significant (all *F >* 114, *p <* 0.001).] In the No Drift condition, similarity with the same pair was very high across training epochs, and similarity was substantially lower in the Drift condition (Figure 7E). This result likely reflected the fact that the input pattern was identical across epochs without drift, whereas it always slightly differed with drift. Intriguingly, however, representational similarity for a given pair against different pairs in the list also markedly differed between the conditions. In this case, similarity to different cues was much lower for the Drift condition throughout training, suggesting there was more separation between patterns in this condition.

### Greater drift between training epochs drives temporal abstraction and decontextualization

In the preceding section, we demonstrated that drift (relative to no drift) benefited long-term memory, produced higher training-related error, and drove pattern separation between memories. Intriguingly, we also found that drift benefited memory when temporal context vectors were scrambled at test, suggesting it improved direct connections between the cues and targets. These results begged the question: does greater drift between learning events benefit memory by strengthening longer and longer time scale representations (temporal abstraction) (Toppino & Gerbier, 2014), does it benefit memory by improving cue-target connections (decontextualization), or both? (Figure 1)

To address this question, we ran simulations with a large range of ISIs and RIs. Specifically, we included 11 ISIs between training epochs [nine Drift conditions with 2 to 512 time steps of drift, plus a No Drift with no drift between learning epochs and a Scrambled ISI condition with a random temporal context vector between each learning epoch] and 9 RIs [seven RIs after the last training epoch, from 32 (2^5^) to 2048 (2^11^) time steps, plus a No Lag condition using the final training temporal context and a Scrambled condition with a random temporal context at test]. We will present the results by speaking generally about relatively short and long ISIs and RIs, and we will summarize and interpret this entire subsection below.

In line with our prior results and canonical spacing effects (Benjamin & Tullis, 2010; Cepeda et al., 2006), the relationship between ISI and RI was non-monotonic and memory was best for small ISI conditions at short RIs (and the No RI Lag condition) and large ISI conditions at long RIs (and the Scrambled RI condition) (Figure 8A). This was supported by a two-way ANOVA on memory performance, with ISI x RI as factors. This showed significant main effects for both factors and a significant interaction (all *F >* 96, *p <* 0.001). Moreover, even in the Scrambled RI condition, longer ISIs produced better recall, suggesting more drift produced more decontextualized, direct cue-target benefits. However, the forgetting effect occurred gradually across RIs, such that the middle ISI condition had better memory recall than the shortest- and longest-ISI conditions by RI-128 (both, *p <* 0.001). Importantly, in the later RIs, such as RI-512, recall was best in the moderate-ISI conditions and still better than in RI-2048 and Scrambled RI conditions (both, *p <* 0.001), suggesting that, in addition to direct cue-target strengthening, there was long-term strengthening of slow-drifting temporal contexts that was eliminated by scrambling the temporal contexts further. This constitutes evidence for long-term temporal abstraction as a process apart from complete decontextualization.

**Figure 8:**
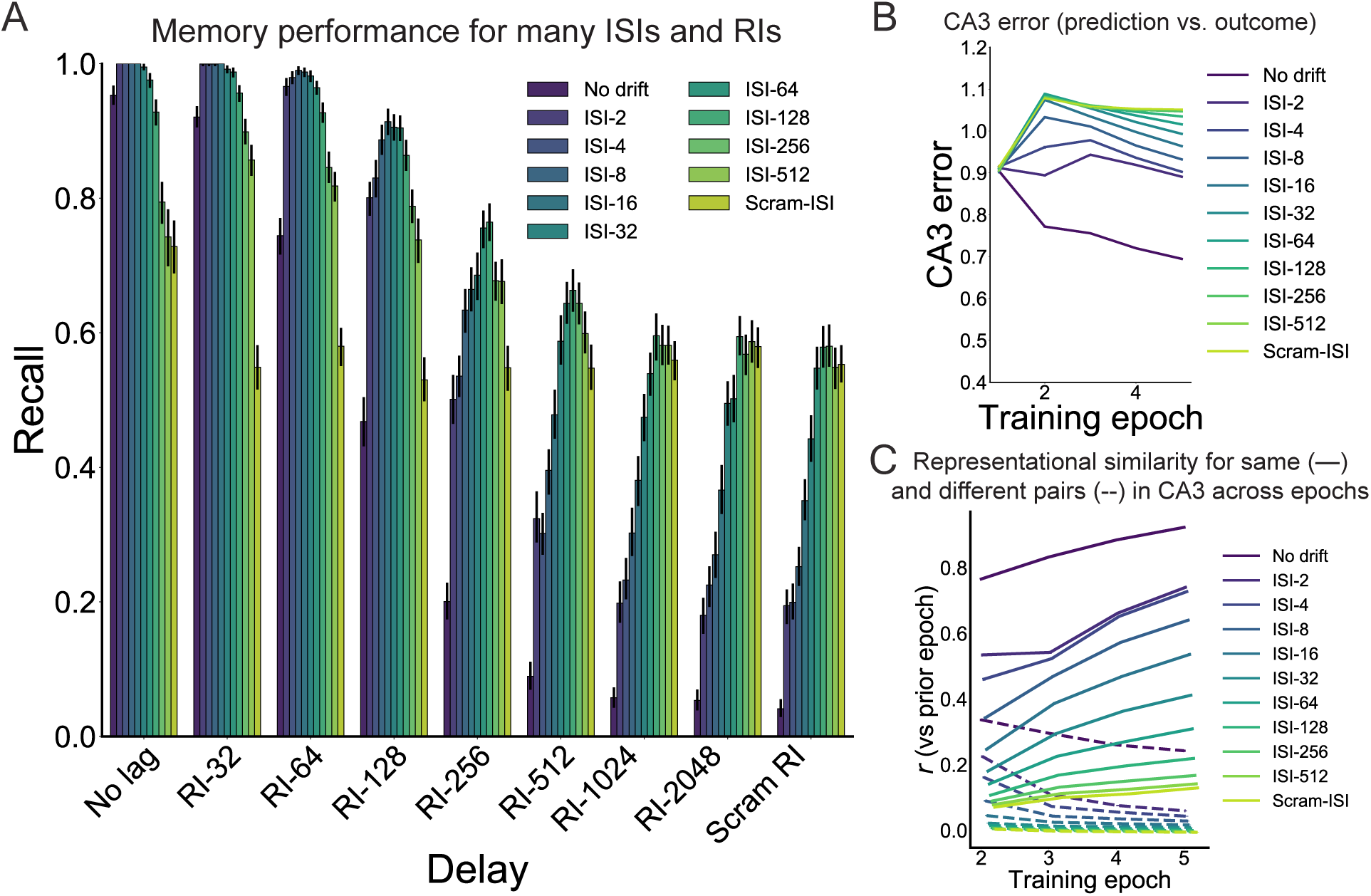
Effects of different amounts of spacing between training epochs on model performance. (A) Different amounts of spacing influenced memory performance depending on the retention interval: shorter spacing benefited memory best at short retention intervals, whereas greater spacing benefited memory best at long retention intervals. (B) Across training, CA3 errors remained higher across epochs with greater spacing. (C) Across training, representations from epoch-to-epoch increased for the same cues (solid lines) and decreased across cues (dotted lines). Greater spacing produced lower same- and across-pair representational stability values.

In accord with findings showing greater CA3 error in the Drift than No Drift condition in the preceding section, we found a main effect of condition [*F*(10,5445) = 1128.6, *p <* 0.001], such that CA3 error increased with ISI (Figure 8B). We also found a main effect of epoch [*F*(4,5445) = 162.6, *p <* 0.001] and a significant interaction [*F*(40,5445) = 160.1, *p <* 0.001].

Additionally, as in the preceding section, we ran a 3-way, ISI x learning epoch (2-5) x pattern status (same vs. different) ANOVA. We again found that all main effects, two-way, and three-way interactions were significant (all *F >* 272, all *p <* 0.001). Notably, greater ISIs produced lower within-pattern similarity, but also lower across-pattern similarity throughout training Figure 8C.

Critically, to address the question of temporal abstraction, we measured the mean temporal context pool weights between ECin *→* CA3 for each ISI against the No Drift condition. A two-way, ISI by temporal context pool interaction on mean weights produced main effects of both factors and a significant interaction (all *F >* 444, all *p <* 0.001). For the interaction, we found a full crossover effect between temporal context pool and ISI: the fast-drifting pools had the greatest mean weights in the short-ISI conditions, whereas the slow-drifting pools had the greatest mean weights in the long-ISI conditions (Figure 9).

**Figure 9:**
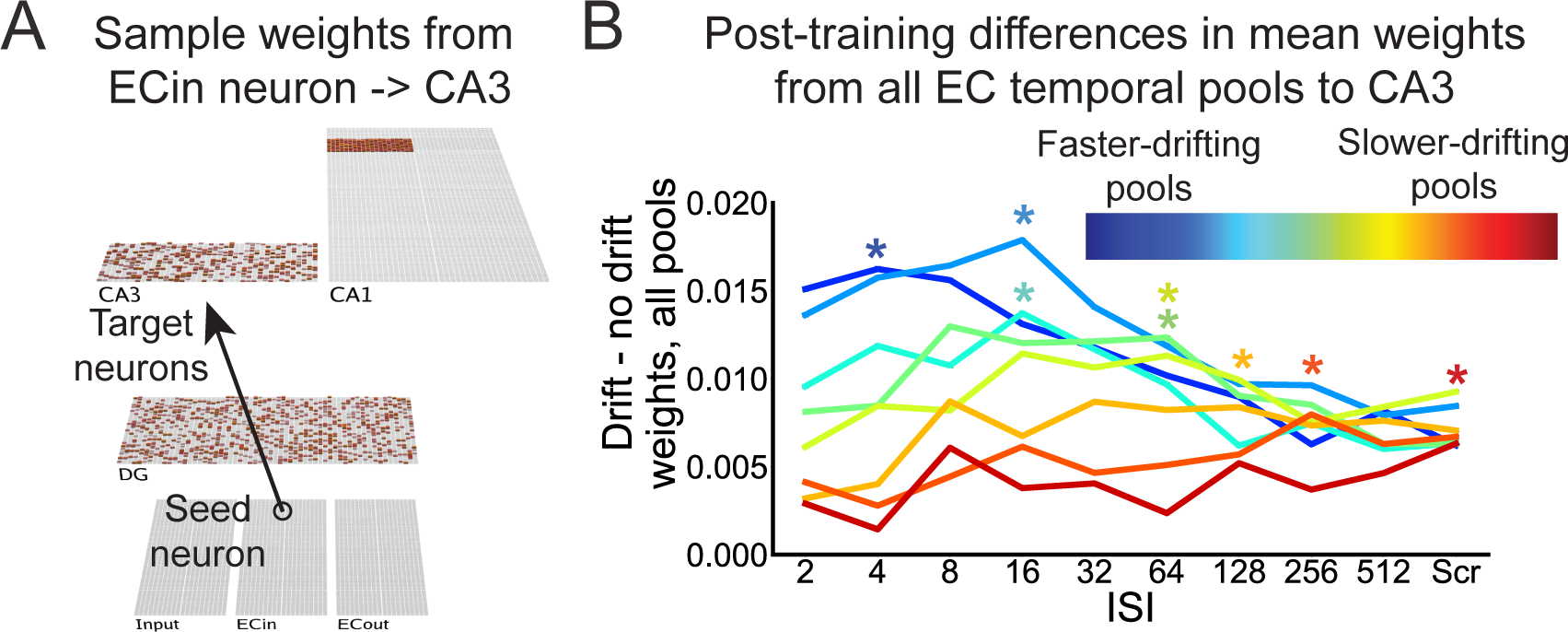
Temporal abstraction in HipSTeR. (A) Weight strengths from a sample seed neuron from ECin *→* CA3 units are shown. (B) Post-training differences in mean weights between each ISI against the No Drift condition are shown for each temporal context pool. Asterisks indicate the ISI with the strongest mean weight for each pool. Across spacing conditions, greater spacing produced weaker weights for the shorter timescale pools and stronger weights for the longer timescale pools.

We now offer a cumulative explanation of these results. The short-RI advantage for short-ISI conditions occurs because the fast-drifting temporal context vectors can still offer cue support to the memory after short RIs, so strengthening these vectors benefits recall at these time points. For short-ISI conditions tested after long RIs, strengthening the fast-drifting temporal context pool weights does very little to support recalling the memory because the test patterns have drifted far away from the final training pattern; this condition has caused the model to effectively overfit to a local temporal context, and therefore forgetting occurs more quickly. The long-ISI conditions are inferior at short RIs because so much drift has occurred between training epochs that, while they can strengthen the slow-drifting temporal context pool weights, they do not strengthen many of the fast-drifting temporal context weights as well; these fast-drifting weights are effectively scrambled by the time new training epochs occur, so each new training instance effectively strengthens a new random subset of weights. Nevertheless, the slow-drifting vectors retain some overlap across training epochs, and their corresponding pool weights become strengthened. Therefore, the slow-drifting pools support memory recall in long-ISI conditions at long RIs better than in short-ISI conditions. This temporal abstraction process confers advantages for the long-ISI conditions at medium-to-long RIs, before the slow-drifting pools have also drifted to a chance level of overlap. Finally, greater error also results in more decontextualized cue-target associations because the same ECin units will be activated for each cue and target, and their corresponding weight changes in HC will be stronger due to the higher error in the network layer. This produces spacing benefits at the longest RIs and when the temporal context pools have become fully scrambled (Figure 1).

### Eliminating learning in specific pathways reveals dissociable learning mechanisms within the hippocampus

We have thus far outlined two primary learning mechanisms that can support spaced learning: temporal abstraction and decontextualization. To examine how different pathways and learning mechanisms in HipSTeR produced these results, we next performed analyses involving turning off learning or specifically EDL in specific pathways. We contrasted the full HipSTeR model in the preceding section against models that were identical except for the following changes: ECin *→* DG (no learning), ECin *→* CA3 (no learning), ECin *→* CA3 (no EDL, but Hebbian learning was present), ECin *→* CA1 (no learning), CA3 *→* CA3 (no learning), and CA3 *→* CA1 (no learning). We will present these results by their increasing relevance for elucidating learning mechanisms.

As expected, turning off learning between CA3 *→* CA1 abolished learning completely in the model, as it is the only connection from the trisynaptic and disynaptic pathways (Figure 10, top center). Conversely, turning off learning from ECin *→* CA1 had almost no effect on performance (Figure 10, top right); however, at later intervals, this model actually outperformed the full HipSTeR model in short-ISI conditions tested at long RIs (more on this below). This was supported by a two-way, ISI x RI ANOVA on memory differences between the control and lesion models, which showed main effects of both factors and, critically, a significant interaction (all *F >* 5, all *p <* 0.001). Turning off learning between the recurrent CA3 *→* CA3 pathway produced moderate impairments relative to the full model that increased with drift (Figure 10, top left), which was supported by a similar ANOVA showing a main effect of RI [*F*(4,780) = 26.7, *p <* 0.001], no main effect of ISI [*F*(3,780) = 0.4, *p* = 0.54] and a significant interaction [*F*(12,780) = 25.3,, all *p <* 0.001].

**Figure 10:**
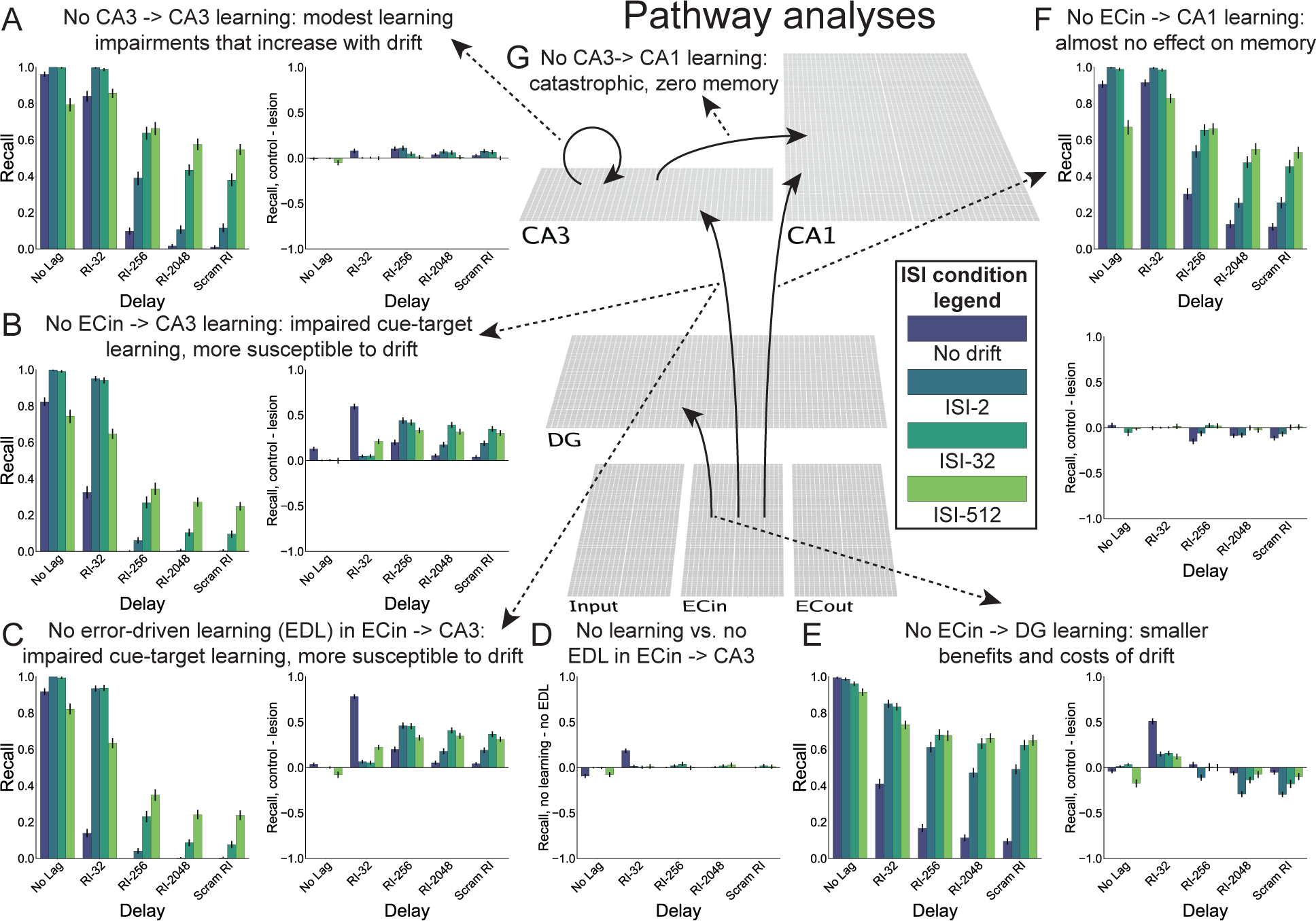
Removing learning in different CLS pathways revealed dissociable learning mechanisms. We plotted HipSTeR model architecture in the center, with affected pathways shown as arrows connecting results on the periphery. For simplicity, we plotted retention interval conditions for only four spacing conditions: No Drift, ISI-2, ISI-32, and ISI-512, representing no, short, medium, and long spacing. (A) Preventing CA3 *→* CA3 learning in the model left performance generally preserved (left). Comparing this with the full HipSTeR model shows only modest improvements for the full model that increase with greater drift (right). (B) Preventing learning from ECin *→* CA3 impaired cue-target learning, making performance highly susceptible to drift (left). Comparisons with the full model showed nearly uniform performance differences across retention intervals (right). (C) Specifically preventing error-driven learning (EDL) in this pathway produced similar results, both on its own (left) and when compared with the full model (right). (D) Directly comparing the full- and Hebbian-only models revealed minimal differences. (E) Preventing learning from ECin *→* DG preserved cue-target learning and created a relative resistance to neural drift: Benefits of the full model are present with short retention intervals but are reversed at longer retention intervals, such that performance is actually superior in this model versus the full model. (F) Preventing learning from ECin *→* CA1 left memory largely intact (top), as shown by negligible differences when comparing it with the full model (bottom). (G) Preventing CA3 *→* CA1 learning abolished memory in all conditions and at all retention intervals (unshown).

We saw the strongest and most interesting dissociation between models without ECin *→* CA3 learning and without ECin *→* DG learning. Turning off learning completely from ECin *→* CA3 strongly affected cue-target learning, such that the memory became hyper-susceptible to drift (that is, performance decreased more quickly with drift) relative to the full HipSTeR model (Figure 10, left middle) [main effect of RI: *F*(4,780) = 230.4, *p <* 0.001; no main effect of ISI:*F*(3,780) = 1.4, *p* = 0.54; interaction: *F*(12,780) = 533.4, all *p <* 0.001]. This was similarly the case when we turned off EDL but kept Hebbian learning in this pathway (Figure 10, bottom left) (both main effects and interaction: *F >* 34, *p <* 0.001); the differences between the complete non-learning and the no EDL ECin *→* CA3 models were significant, but they were quantitatively small (Figure 10, bottom center) (both main effects and interaction: *F >* 142, *p <* 0.001). These results suggest that EDL from ECin *→* CA3 strongly drives the drift-resistant, temporally abstracted or decontextualized part of the memory directly linking cue and target. Critically, they also reveal a novel mechanism by which decontextualization – generally thought to be a process confined to the neocortex (Hasselmo, 2005; Winocur, Moscovitch, & Bontempi, 2010) – could occur within the hippocampus itself.

As opposed to the ECin *→* CA3 pathway, turning off ECin *→* DG learning rendered the memory largely resistant to drift. Relative to the full HipSTeR model, this model mostly lacked the benefits conferred by the intact temporal context at short RIs, which have temporal contexts that have not yet drifted away from their training contexts (Figure 10, bottom right) (both main effects and interaction: *F >* 26, *p <* 0.001). On the other hand, without learning in this pathway, error in CA3 remained higher during training, which helped to strengthen the direct cue-target aspect of the memory (dependent on ECin *→* CA3). As a result, this model paradoxically outperformed the full HipSTeR model at long RIs. (The very weak benefits at long RIs in the non-learning ECin *→* CA1 model likely occurred for similar reasons.) In sum, the more direct ECin *→* CA3 pathway produced the spacing benefits of temporal abstraction and decontextualization that make memories more drift-resistant, whereas the ECin *→* DG pathway produced transitory benefits that continued to rely on temporal context and the scale of which depended on the amount of drift during training.

### Comparing multi-scale to uniform drift

Many prior temporal context models posit that drift occurs across multiple time scales [e.g., Liu et al. (2019)], which is supported by various forms of neurobiological evidence [e.g., Tsao et al. (2018); Umbach et al. (2020)]. Nevertheless, it is worthwhile to assess the advantages and disadvantages of a system employing uniform as opposed to multi-scale drift. In the next three simulations, we contrasted our results across numerous ISIs and RIs from our multi-scale (control) model against identical models using uniform drift for all eight temporal context pools. We chose three drift rates that were equal to the fastest, slowest, and a medium-speed pool from our control model, and we plot the results alone and against our control model (Figure 11). For statistics, we ran two-way, ISI x RI ANOVAs on the memory differences for each model against the control model. First, for most ISIs and RIs in the all-fast drift model, the pattern is completely new upon each training instance, because the auto-correlated pattern reaches a value of 0 after approximately 16 time steps (Figure 11A) (both main effects and interaction: *F >* 18, *p <* 0.001). This means that the amount of drift is a much smaller factor, so most of the results after the No Lag RI are equivalent (Figure 11B). With 5 training epochs, the model actually outperforms the multi-scale model after modest-to-long delays because the high error (from having essentially random context patterns on each training instance) drives such strong decontextualization that the memories are robust to drift. However, at short delays like RI-32, the multi-scale model outperforms the all-fast model due to lack of cue support in the all-fast model.

**Figure 11:**
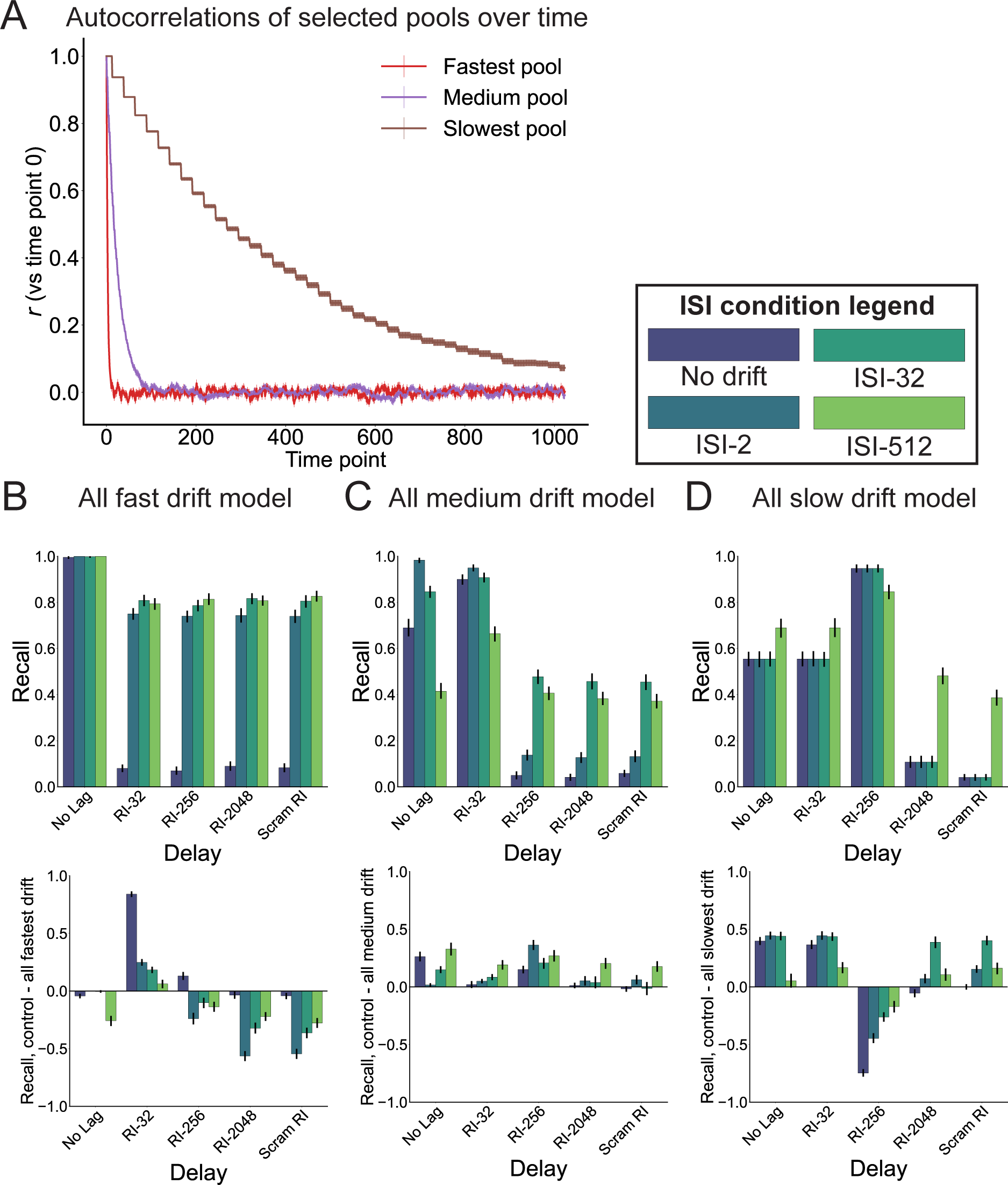
Simulations contrasting multi-scale drift against uniform drift. (A) Autocorrelations for the three uniform drift rates (equivalent to Pool 1, 5, and 8 in the multi-scale model, respectively). (B-D) Results from models with the fastest, medium, and slowest uniform drift rates (top) and direct contrasts between the multi-scale (control) model against these models (bottom).

Next, we will analyze the all-slow model (Figure 11D). In contrast to the all-fast model, after many ISIs and RIs, the temporal context pattern is almost completely intact (both main effects and interaction: *F >* 20, *p <* 0.001). When the ISI is low (No drift or short ISI conditions, such as ISI-32), all cue-target pairs from the 5 epochs are trained on almost exactly the same temporal context pattern. Note that this differs somewhat from the ‘No Drift’ condition in prior simulations, in that the drift here is so slow that there is hardly any within-epoch drift in addition to little between-epoch drift at short ISIs. This means that the retrieval context after a short delay (such as with no lag or at RI-32) will resemble the pattern of the final training epoch – and there will be massive interference among list items. Interestingly, the model actually benefits from some drift to “unstick” the memory from having basically the same pattern across all epochs, which, when it remains the same for all pairs within the list, causes interference among the pairs. These performance benefits arise with either moderate RIs (e.g., RI-256), which are long enough to prevent some of the unsticking that occurs at low RIs but not so long that the very slow-drifting patterns nonetheless drift away (e.g., RI-2048). The benefits can also occur with more drift during learning epochs (e.g., ISI-512). In these cases, the model learns on different temporal patterns on each instance and does not repeatedly strengthen the same (ultimately interfering) weights. The model is inferior to the multi-scale model at most intervals because of either this interference (at short RIs) or reduced error-driven learning because the pattern remains intact for longer (at long RIs). However, it is superior to the multi-scale model at a moderate RI like RI-256, indicating that there is some retention interval for which it is optimized. Ultimately, the slow drift model allows patterns to be maintained for long periods of time but suffers from interference due to poor within-list drift. Finally, the all-medium model falls between these two extremes, performing closest to the control model of the three simulations but showing deficits with very short or long lags (Figure 11C) (both main effects and interaction: *F >* 17, *p <* 0.001).

It is never certain during learning when information will be re-learned or need to be retrieved. Therefore, we interpret these results to indicate that multi-scale drift does not optimize for any particular RI but rather a range of RIs, balancing learning and long-term memory maintenance while allowing for re-learning benefits according to the temporal frequency of the information.

### Decontextualization in other paradigms

To the best of our knowledge, conceptualizing spacing effects as temporal decontextualization is novel. Therefore, our final simulations aimed to bridge our modeling framework to other studies falling under the umbrella of decontextualization, which have used paradigms with environmental, task, or background (pictorial) contexts (Butler, Black-Maier, Raley, & Marsh, 2017; Glass, 2009; Maskarinec & Thompson, 1976; S. M. Smith et al., 1978; S. M. Smith & Handy, 2014, 2016; S. M. Smith & Rothkopf, 1984; Soderstrom & Bjork, 2015; Trask & Bouton, 2018; Zawadzka, Baloro, Wells, Wilding, & Hanczakowski, 2021). These final simulations thus demonstrate that our general modeling approach can also capture other decontextualization effects.

In decontextualization paradigms, the contexts are either constant or variable before final tests take place in novel contexts. In S. M. Smith et al. (1978), subjects learned words in room A and either practiced recalling the words in room A or in room B in a second session before taking a final test in room C. Critically, memory performance in the variable room (relative to the constant room) was worse during practice but better at the final test in room C, suggesting the environmental variability decontextualized the memory. Imundo et al. (2021) used a similar method but had subjects either re-study or retrieve lists (without feedback) during the second session. They found variability benefits, but only during restudy, whereas retrieval (without feedback) likely relied on successful retrieval for benefits. In S. M. Smith and Handy (2016), subjects learned word pairs against background contexts that were unique for each pair and either remained constant across five practice trials or varied across trials. A final test was given for all pairs with no background context. As in S. M. Smith et al. (1978) and Imundo et al. (2021), cued recall for pairs in the variable (relative to the constant) condition was worse during learning but better at the final test, suggesting trial-specific pictorial context variability also decontextualized memories. Following the logic outlined in the preceding sections, these findings could arise because variable contexts offered less cue support to memories during learning, and this therefore led to more direct strengthening of the cue-target elements that were in common across training epochs. These changes would then support the memory when tested in novel contexts. Note that, for the purposes of our simulations, S. M. Smith et al. (1978) manipulated context at the epoch level, whereas S. M. Smith and Handy (2016) manipulated context at the trial pair level. Note also that if the test in these paradigms occurred instead with the original training context, pairs in the the constant (relative to the variable) context should produce better memory, in line with some of both their results (S. M. Smith et al., 1978) and findings akin to context-dependent memory (Godden & Baddeley, 1975). These effects should arise because weights related to the original training context should be repeatedly strengthened across training (Cox, Dobbelaar, Meeter, Kindt, & van Ast, 2021; Estes, 1955a; S. M. Smith & Handy, 2014, 2016).

To simulate these results, we kept the same HipSTeR model architecture and slightly modified the inputs (Figure 12A). The inputs included the same number of cue and target pools and the six fastest-drifting temporal context pools (out of eight). Here, instead of the two slowest drifting temporal context pools, we added two other context pools. In our epoch-wise simulations (representing environmental context, as in S. M. Smith et al. (1978)), the context vectors were identical throughout a learning epoch. Across epochs, they either remained the same (Constant) or changed for each epoch (Variable). At final test, these pools had either the context vectors from the first learning epoch (Old) or a new vector (New). In our trial-wise simulations [representing background pictorial contexts, as in S. M. Smith and Handy (2016)], the context vectors were unique for each pair. Across epochs, they either remained the same for that specific pair (Constant) or changed on each epoch (Variable). At final test, these pools had either the same specific context vectors for that pair from the first learning epoch (Old) or a new vector (New). In all experiments, we used a temporal ISI of 2 steps and RI of 512 steps.

**Figure 12:**
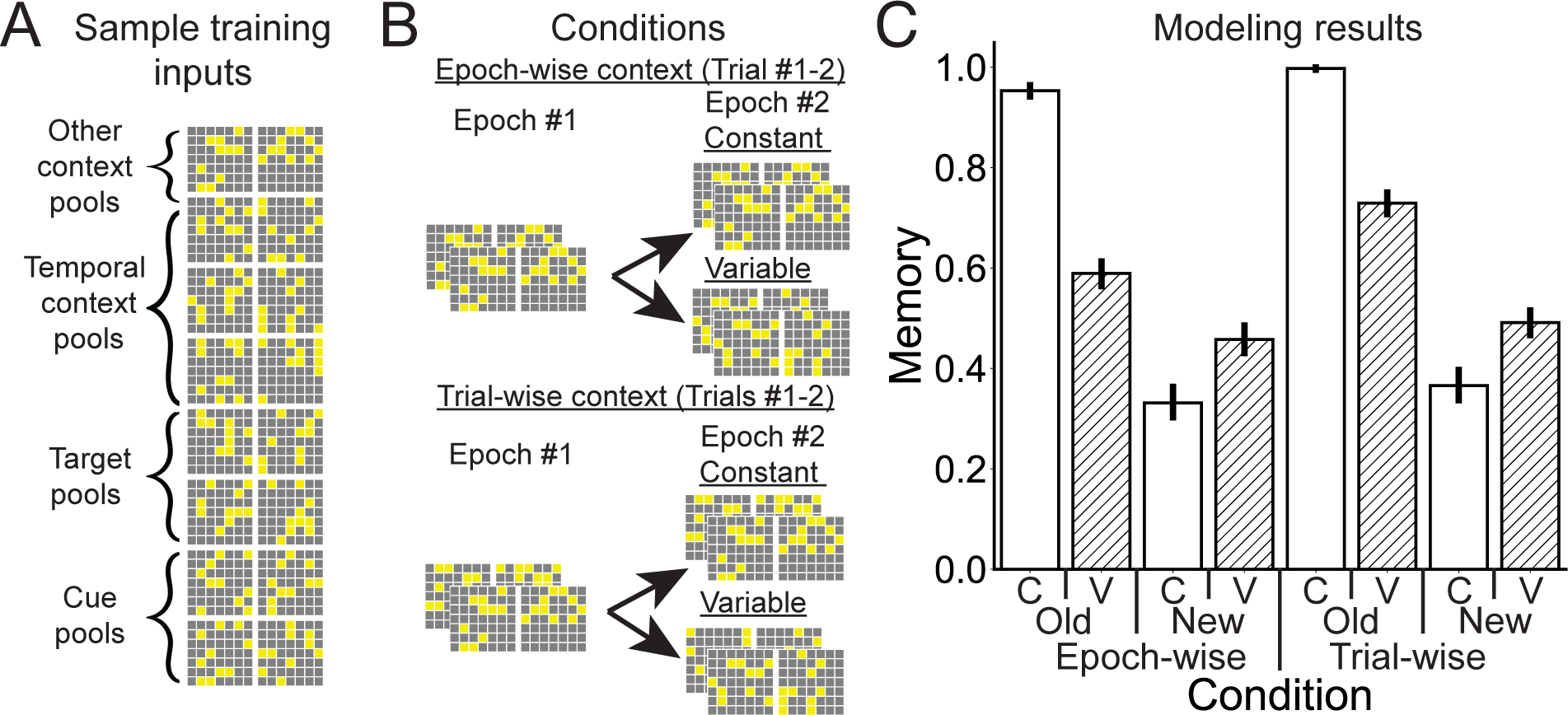
Simulations of decontextualization in other paradigms. (A) For these simulations, we modified HipSTeR model inputs slightly by replacing two temporal context pools with other context pools. (B) Other context pools were either constant within an epoch (top) or unique for each cue-target pair (bottom). (top) Epoch-wise contextual changes were always constant within an epoch (top left) and either constant across all epochs or variable across epochs (top right). (bottom) Trial-wise contextual changes were always different for every trial even within an epoch (bottom left) and were either in a constant order in each epoch or variable for each trial (bottom right). (C) Modeling results showed that, while performance was superior in the constant training context when the constant (old) context was given at test, performance was superior in the variable training context when a new context was given at test. C: constant training context, V: variable training context.

We analyzed these data with a three-way, training type (epoch-wise vs. trial-wise) x test context (old vs. new) x training variability (constant vs. variable) ANOVA on model memory. We found that all three main effects, all three two-way interactions, and the three-way interaction were all significant (all *F >* 40, *p <* 0.001). The main effect of training type showed that overall memory performance was higher in the trial-wise than epoch-wise simulations. This likely occurred because the more specific context cues resulted in more efficient learning with less incidental interference resulting from the same context applied to all pairs of a list (Nairne, 2002) (Figure 12B). The other main effects indicated that memory was better overall with constant training and old contexts presented at text, but note that the interactions are of more interest here. In line with prior results and our predictions, in both epoch-wise and trial-wise simulations, final test performance was superior in the constant relative to the variable condition when tested with the old context (*p <* 0.001), showing unsurprisingly that cue support from training aids memory (Cox et al., 2021). More critically, however, performance was superior in the variable relative to the constant condition when tested with a new context (*p <* 0.001), suggesting that contextual variability led to decontextualized memory traces that allowed for successful memory recall even with new contexts.

## Discussion

By implementing drift and EDL within HipSTeR, a biologically plausible model of the EC-HC network, we simulated forgetting and a wide array of spacing effects from the cognitive psychology literature. First, we ran simulations that replicated specific spacing effect principles. We found that the optimal spacing interstimulus interval (ISI) depended on the retention interval (RI) before the final test (Cepeda et al., 2008), that relearning after long temporal delays overrode smaller differences due to initial spacing (Rawson et al., 2018), that different spacing schedules produced modest differences (Küpper-Tetzel et al., 2014) but that absolute spacing was the most critical factor (Karpicke & Bauernschmidt, 2011), and that spacing can produce direct benefits between cues and targets at extremely long RIs – intervals when there is little (to arguably no) resemblance between the temporal contexts during training and at test [as in the 5-year followup test in Bahrick et al. (1993)].

Next, we probed the mechanisms producing these effects in HipSTeR. The very presence of temporal context drift (relative to no drift) resulted in better recall performance, greater error in CA3 in later training epochs, and greater representational dissimilarity between different CA3 activation patterns. Further analyses investigating a full spectrum of drift values between training epochs showed that greater spacing produced more temporal abstraction (as shown via stronger mean weights in slower temporal context pools from ECin *→* CA3) and decontextualization (as shown via better recall when temporal context vectors were scrambled) (Figure 1). These analyses also showed that massed learning can be superior in the short term because there is preferential strengthening in the faster drifting temporal context vectors, which can still provide cue support for the memory after low drift.

Turning off learning in various pathways in HipSTeR showed a stark dissociation between a pathway that continued to support contextualized memories and benefited the model under conditions of low drift between the training and testing context (ECin *→* DG *→* CA3) and one that supports decontextualization (ECin *→* DG *→* CA3). The latter mechanism is especially novel because decontextualization has generally been attributed to the neocortex (Winocur et al., 2010; Yassa & Reagh, 2013), but here we show how it can arise in the hippocampus. Comparing our model using multi-scale drift against a set of uniformly drifting alternatives showed that, while the uniform models performed better when testing after some particular RIs, the multi-scale model seemed to balance memory performance for a range of potential RIs. Finally, because conceptualizing spacing effects as arising partly due to decontextualization is novel, we aimed to link our results with more canonical decontextualization effects in the literature. Using an identical model architecture and similar inputs, HipSTeR captured classic decontextualization effects showing that variable encoding contexts lead to better memory when tested in novel contexts.

### Comparisons with other spacing effect and encoding variability theories and models

Many papers argue that spacing effects could arise from some combination of two processes: encoding variability, which determines the amount of strengthening, and study-phase retrieval, which determines the likelihood of strengthening (Benjamin & Tullis, 2010; Küpper-Tetzel & Erdfelder, 2012; Landauer, 1969; Mozer et al., 2009; Raaijmakers, 2003; Smolen, Zhang, & Byrne, 2016; Walsh et al., 2018). Encoding variability theory asserts that re-encoding benefits increase with more variable encoding contexts (such as after greater temporal lags) because there will be a greater diversity of contextual elements added to the memory, resulting in more routes to retrieval (Estes, 1955a; Maddox, 2016; Mozer et al., 2009; S. M. Smith et al., 1978; S. M. Smith & Handy, 2014). Study-phase retrieval asserts that in order to benefit from spacing, subjects must retrieve the episode from encoding (Thios & D’Agostino, 1976). Ultimately, because encoding variability would produce benefits that increase monotonically with spacing and because study-phase retrieval would be best immediately after encoding and therefore would produce anti-spacing benefits as a solo mechanism, the two have been proposed in combination to capture the non-monotonic nature of spacing effects (Benjamin & Tullis, 2010). From these foundations, a question arises as to what representations become strengthened and why. One intriguing theory is that there is a temporal abstraction process that depends on the ISI (Mozer et al., 2009; Toppino & Gerbier, 2014). That is, longer ISIs may preferentially strengthen aspects of the memory trace that have more long-term stability. In particular, Mozer et al. (2009) created a multi-temporal scale neural network model in which strengthening preferentially occurred in longer-term representations as a function of the error between encoding contexts along shorter time scales, such that a failure to support the memory along shorter time scales forced strengthening along a longer time scale.

How do these theories and models square with results from HipSTeR? We found that encoding variability (in the form of temporal context vector differences) benefited memory at the longest RIs via two EDL-driven mechanisms (Figure 1). Like Mozer et al. (2009), greater ISIs caused temporal abstraction, or greater relative strengthening in slower-drifting temporal pool weights. Greater ISIs also allowed for better intact memory in the face of the complete decontextualization of fully scrambled context vectors. There is therefore a critical difference between how variability produces strengthening in encoding variability theory and in HipSTeR: encoding variability theory suggests that strengthening occurs because the same memory accrues more contextual routes to retrieval, whereas our model suggests it occurs because greater error produces strengthening of the relevant temporal context pool weights and/or cue-target weights. That is, our results suggest that gaining more unspecified contextual routes to the memory may not be ultimately that helpful, since it is unclear, given that drift will occur randomly, how or why any of them should be meaningfully activated during retrieval after further drift has occurred.

Instead, drift in HipSTeR strengthened the weights of slower drifting units that were more likely to be active at retrieval or decontextualized cue-target weights that were fully drift-resistant. We believe this latter explanation seems almost necessary to explain results at extremely long RIs, such as the 5-year RI from Bahrick et al. (1993) or results in other paradigms showing retention across decades (Maxcey, Shiffrin, Cousineau, & Atkinson, 2022). Moreover, this account can be considered even more straightforwardly in our final environmental/pictorial decontextualization paradigm simulations. We tested these models with novel context vectors, and there was no reason why these (random) vectors should have activated any of the contextual elements accrued in the variable condition any more than they would have activated unhelpful elements that would have counteracted memory retrieval. Altogether, rather than encoding variability driving memory strengthening while study-phase retrieval limits the likelihood of strengthening, in HipSTeR, EDL drove strengthening via temporal abstraction and decontextualization while contextual drift produced forgetting with elapsing time.

A recent paper showed that as rodents accumulated experience within the same environment across multiple days, some place cells – or cells that fire at particular locations within a spatial context – transiently entered and left the memory trace, whereas other ones sustained a stable firing location across days (Vaidya, Chitwood, & Magee, 2023). Intriguingly, the proportion of sustained neurons grew over time and predicted behavioral performance, in accordance with the memory trace reactivating and solidifying itself with experience. We believe that an experiment manipulating spacing and using a similar analytical approach that also measures their endurance (how long they stay in the memory trace) could help adjudicate between the predictions of encoding variability and our model. That is, encoding variability theory predicts that spacing would increase the *proportion* of new neurons in a trace upon relearning (even if they were ultimately transient and not part of the trace in the next learning session). Conversely, our model predicts that spacing would increase the number and endurance of sustained neurons in the trace. A further prediction of our model, given the principle of temporal abstraction, is that the amount of spacing should directly scale with neuron endurance.

These findings resonate with an influential theory that separates memory constructs into retrieval and storage strength (R. Bjork & Bjork, 1992; R. A. Bjork, 2011). Retrieval strength refers to the in-the-moment accessibility of a memory and explains memory performance at a particular time, whereas storage strength refers to a latent factor referring to how well the memory is learned and explains its persistence at later moments. [Note that prior researchers similarly distinguished between constructs related to current and later performance (Estes, 1955a, 1955b; Hull, 1943; Skinner, 1938).] Mapping these constructs onto our modeling findings, we believe retrieval strength relies on momentary pattern matches that depend partially on the overlap between the learned temporal context and the temporal context at retrieval. Conversely, both temporal abstracted weights (which help the memory for longer periods of time but not necessarily forever) and decontextualized weights (which directly strengthen cue-target associations) align with the concept of storage strength.

### The organization of time and model plausibility

Researchers have long been interested in the role of time in episodic memory. “Mental time travel” was a core aspect of Tulving (1972)’s conceptualization of episodic memory, suggesting the reinstatement of a particular temporal frame. Countless researchers have investigated the organization of time in memory, with a core question being whether or not it is organized by chronological “time stamps” (Bradburn, Rips, & Shevell, 1987; Burt, 1992; Friedman, 1993; Hintzman, 2016; Jeunehomme, Folville, Stawarczyk, Van der Linden, & D’Argembeau, 2018). Indeed, contiguity effects in memory, whereby information presented nearby in time are also temporally clustered together during recall, suggest that individuals reinstate temporal contexts that guide further retrieval [e.g., Howard and Kahana (2002); L. J. Lohnas et al. (2015)]. Intriguingly, temporal clustering occurs across numerous time scales, across lists, and up to days and months Healey et al. (2019); Howard et al. (2008); Moreton and Ward (2010); Uitvlugt and Healey (2019); Unsworth (2008), suggesting a scale-invariant property in support of log-spaced temporal representations (Brown et al., 2007; Howard, 2018). Moreover, reinstatement has also shown neural contiguity effects (Folkerts et al., 2018; Manning et al., 2011), and it has been proposed that another role of neural drift is to create such time stamps (A. Rubin et al., 2015).

However, there have been notable criticisms of these ideas [e.g., Friedman (1993); Hintzman (2016)]. Many free recall tasks rely on intentional encoding, whereby subjects can develop rehearsal strategies that rely on successive rehearsals, and incidental encoding drastically reduces (though does not eliminate) temporal contiguity effects (Dester, Lazarus, Uitvlugt, & Healey, 2021; Healey, 2018; Mundorf, Lazarus, Uitvlugt, & Healey, 2021). Moreover, temporal contiguity effects can be reduced or altered by other types of (e.g., semantic, narrative) structure [e.g., Antony et al. (2021); Bousfield (1953); Heusser, Fitzpatrick, and Manning (2021); Polyn et al. (2009)], suggesting these effects are highly manipulable in the presence of more salient organizing characteristics. Additionally, temporal context theories might predict that successively presented cued recall trials would differ as a function of their lag at encoding, but multiple experiments have shown no effect of this lag (Osth & Fox, 2019). At the very least, these findings question the idea that we automatically encode and retrieve information as if reading from a timeline.

How can we resolve these ideas and determine to what extent drift might influence memory? We propose that time is a weak signal if not given prioritized attention. That is, the ability to reconstruct the past in a direct linear order when one is not intending to or does not encode a set of causally connected events in a linear order is poor (Dimsdale-Zucker et al., 2020), but not zero (Dester et al., 2021; Healey, 2018). Even in a rodent experiment that showed robust decoding of trial number from LEC neurons, when a task change eliminated the need to keep track of different trials, the ability to decode trial number from the neurons was reduced, but not eliminated [Figure 8 experiments, Tsao et al. (2018)]. It is reasonable then to wonder what the point of such a drifting signal could be across each of these time scales. We speculate that – just like features of the environment that are represented in sensory subsystems but do not receive direct attention – the ability to use this signal relies strongly on whether one incorporates it somehow into the focus of attention (Niv et al., 2015). Though speculative, this may even be optimal from a memory standpoint, as too strong of a temporal signal may result in linking unlike events that *just so happen* to co-occur. This idea suggests that the prominence of drifting signals within a memory trace can fall under executive control, either becoming accentuated or minimized [see below Limitation on neural drift versus shift; DuBrow et al. (2017)].

### Relevance to temporal context models of memory

Dating back to Estes (1955a) and Bower (1967), memory models have considered context as a pattern of activity that fluctuates over time. This family of models has shown that modeling context can capture numerous temporal memory phenomenon. For instance, Mensink and Raaijmakers (1988) captured how memory interference depends on the timing of interfering information and the time of test, capturing intricate findings from the literatures on proactive and retroactive interference and spontaneous recovery. Later, Howard and Kahana (2002) showed how binding learned items to a drifting context can capture aforementioned temporal contiguity effects in free recall. These models have also generalized this idea to other types of context, such as semantic (L. J. Lohnas et al., 2015; Polyn et al., 2009) and emotional context (Talmi, Lohnas, & Daw, 2019). Altogether, this family of models has simulated an impressive array of memory data.

The primary contribution of our model to these efforts is to show how memories get updated and strengthened upon repetition. Context continually drifts over time in a multi-scale manner, but our model suggests that repeating information at increasing timescales will strengthen increasingly long context representations that effectively help the model generalize over time. Therefore, in HipSTeR, memory strength at a given time relies on a rich combination of drifting temporal context and the timing history of training.

### Implications for the neurobiology of drift

The theory behind HipSTeR draws heavily from recent neurobiological evidence of drift in single neurons and neural ensembles within the MTL. This evidence stems from two types of findings. The first involves activity from individual neurons or ensembles that ramp up or down spontaneously (Folkerts et al., 2018; Howard et al., 2012; Tsao et al., 2018; Umbach et al., 2020; Yoo, Umbach, & Lega, 2022) or in a cue- or context-evoked fashion (Bright et al., 2020; Liu et al., 2022; Tsao et al., 2018; Umbach et al., 2020; Yoo et al., 2022). This “ramping” drift, which constitutes a form of persistent activity, could occur as a result of unique, slow-adapting Ca++ currents (Egorov, Hamam, Fransén, Hasselmo, & Alonso, 2002; Liu et al., 2019; Tahvildari, Fransén, Alonso, & Hasselmo, 2007; Tiganj et al., 2015). The second compares representational similarity between successive experiences of the same type after variable delays, such as neural activity patterns when animals are placed in the same environment (Bladon, Sheehan, De Freitas, & Howard, 2019; Keinath & Brandon, 2022; Mankin et al., 2015; Marks & Goard, 2021; Mau et al., 2018; A. Rubin et al., 2015; Rule, O’Leary, & Harvey, 2019; Y. Ziv et al., 2013). This drift has been attributed to fluctuations in synaptic weights (Mau et al., 2020; N. E. Ziv & Brenner, 2018) and intrinsic excitability (Delamare, Zaki, Cai, & Clopath, 2023), and the difference in these drift types may be critical when considering temporal effects between those from seconds to hours against those from days to years. Therefore, although HipSTeR implements drift in a manner more resembling the former, ramping type, it is worth considering in later models whether these results would hold in models with drifting synaptic weights.

One common puzzle arising within this literature on drift is how long-term memories survive amidst drift and what function drift might serve. It has been claimed (and shown via modeling) that long-term representations can continue to survive even with substantial drift (Kalle Kossio, Goedeke, Klos, & Memmesheimer, 2021; Mau et al., 2020; Rule & O’Leary, 2022). Using a recurrent neural network model, Clopath, Bonhoeffer, Hübener, and Rose (2017) showed that long-term stability can be maintained with a “backbone” of stable neurons and recurrent activity. Relatedly, while there may be drift from the perspective of some external input (e.g., the environment), internal representations between neural representations and their downstream readers may remain relatively coherent and low-dimensional latent structure relatively constant, which could help to compensate for drift in earlier representations (Delamare et al., 2023; Gallego, Perich, Chowdhury, Solla, & Miller, 2020; Kalle Kossio et al., 2021; Mau et al., 2020; Rule et al., 2019).

Drift could also be helpful for new encoding, as it allows for greater flexibility and even forgetting of old information in an arguably rational manner. Regarding flexibility, drift allows efficient new encoding in a way that does not fully depend on or compete with old memories (Frank et al., 2018; Mau et al., 2020), as it allows for new memories to not simply be reassigned to the same, most excitable units (O’Reilly, Wyatte, & Rohrlich, 2017; Rogerson et al., 2014; Zhou et al., 2009). This would allow memories to form with sparse, efficient representations while eventually (after enough time) involving most or all of the relevant neurons in a particular region. Regarding forgetting from a rational memory standpoint, a memory encountered once or a series of experiences repeated within a short temporal frame is less likely to be relevant for the long term than those repeated more infrequently (Anderson & Milson, 1989; Anderson & Schooler, 1991; Mozer et al., 2009). That is, the likelihood of many stimuli in the environment being repeated falls off rapidly as a function of its last occurrence, a principle which has applications in such wide-ranging domains as word occurrences within *New York Times* articles and utterances from parents to children (Anderson & Schooler, 1991). Given computational constraints, it would seem rational to forget such memories to maintain both the potential to form new memories or preferentially retain those more likely to serve future needs on a timescale that resembles the frequency of encoding (see Figure 11). We therefore propose a novel mechanism of drift, such that it allows for useful forgetting and, when a memory becomes experienced (or reactivated) after variable delays, it provides for the potential for strengthening on a relevant timescale in the form of greater EDL-driven temporal abstraction. Additionally, to the extent that drift between training epochs constitutes a form of noise between input patterns, these results – whereby drift helps the network generalize over time – support the idea that optimal levels of noise help avoid overfitting and improve generalization in neural networks (Elman & Zipser, 1988; Hinton & van Camp, 1993; Sietsma & Dow, 1991; Srivastava, Hinton, Krizhevsky, Sutskever, & Salakhutdinov, 2014; Tran et al., 2022).

### Relationship to theories of hippocampal contributions to episodic memory

Our results have relevance for findings relating hippocampal activity to memory of different ages. First, univariate HC activity measured using fMRI increases when recalling memories with increasing temporal lags across trials (Brozinsky, Yonelinas, Kroll, & Ranganath, 2005) or days (Chen et al., 2016). Moreover, the rate of hippocampal ripples increases with the recall of older autobiographical memories (Y. Norman, Raccah, Liu, Parvizi, & Malach, 2021). In HipSTeR, this increase could be spurred by the amount of error encountered between the difference in temporal contexts. Second, a neural version of the encoding variability account – whereby more variability in neural representations across repetitions predicts better long-term memory – has received support from at least two fMRI studies investigating representational similarity in the hippocampus. That is, representational variability (or instability) in hippocampal patterns across repetitions promotes subsequent memory (Karlsson Wirebring et al., 2015) and memory updating (Speer, Ibrahim, Schiller, & Delgado, 2021). These results differ markedly from similar investigations in the neocortex, where it has been found instead that stable patterns across repetitions of the same stimuli promote memory (Xue et al., 2010). One proposed resolution to these disparate findings is that cortical fidelity is required to represent the content of memories, and thereby ensures proper reactivation of old traces, but variability within the hippocampus promotes greater subsequent updating (Karlsson Wirebring et al., 2015). This interpretation fits well with our account, which requires stable inputs about informational content of cues and targets from elsewhere in the cortex into the EC-HC system but also needs variability from temporal context drift to drive optimal learning via EDL. One potentially conflicting result comes from an important recent study showing that the extent to which prior encoding patterns of picture stimuli are reinstated in the hippocampus during re-encoding days to months later predicts subsequent temporal memory for when the picture was first shown (Zou et al., 2023). It is unclear whether this result, whereby pattern stability rather than variability benefits memory, is specific to the temporal memory task, whether there is some other form of undetected neural variability that serves as a strengthening mechanism, or whether it is problematic for the proposed account.

Our results also have relevance for learning within specific HC pathways. The trisynaptic route involving DG formed associations that continued to change and rely on temporal context with drift between training epochs (Pereira et al., 2007), which allowed for multiple traces of a memory that otherwise had identical content like the cue and target (Guzman et al., 2021). This finding aligns with the idea that the drifting excitability shown in DG neurons may be responsible for time encoding (Aimone, Wiles, & Gage, 2006, 2009). Intriguingly, we found the EC *→* CA3 pathway to be critical for the drift-resistant memory component. A prior computational model suggested that this pathway is especially important for generalizing over a variable set of examples belonging to the same visual category at retrieval (Kowadlo et al., 2019). Like our results, the Kowadlo et al. (2019) generalization effects were similarly much stronger for this pathway than for the CA3 *→* CA3 recurrent collaterals, suggesting a prominent role in the EC *→* CA3 pathway at retrieval. In accordance with these results, our effects can be thought of as creating representations that generalize across time, avoiding the pitfalls of memory that occur without any temporal context support.

Due to the profound long-term benefits of spacing, it is natural to ask whether spacing promotes systems consolidation, or the strengthening of the more relatively long-lasting synapses in the neocortex and the gradual ability to recall memories independent of the hippocampus (Attardo, Fitzgerald, & Schnitzer, 2015; Carpenter, 2020; C. D. Smith & Scarf, 2017). However, current evidence on this is unclear. There is no current neuroimaging evidence to our knowledge that suggests that spacing decreases hippocampal involvement. Rather, following spaced versus massed learning, the HC shows increased activity (Li & Yang, 2020; Nonaka et al., 2017) or greater connectivity with neocortex (Ezzyat, Inhoff, & Davachi, 2018) at retrieval. Evidence from amnesics on the necessity of the hippocampus for spacing effects is also inconclusive. Amnesics show no spacing benefits for recollection, a process which requires the hippocampus, but intact spacing benefits for familiarity (Verfaellie, Rajaram, Fossum, & Williams, 2008), which can rely on neocortical sources (Yonelinas, 2002). Developmental amnesics do show spacing benefits in free recall, but their hippocampal size was only diminished by 30%, indicating that benefits could result from residual hippocampal tissue (Green, Weston, Wiseheart, & Rosenbaum, 2014). Our results suggest that spacing benefits could be explained without systems consolidation. However, we hope to test the impact of the cortex in later models, especially given evidence that a related phenomenon known as the testing effect – whereby long-term memory benefits more from retrieval than re-study – results in greater neocortical involvement (Antony, Ferreira, Norman, & Wimber, 2017; Ferreira, Charest, & Wimber, 2019; Himmer, Schönauer, Heib, Schabus, & Gais, 2019; Van den Broek, Takashima, Segers, Fernández, & Verhoeven, 2013). One particularly interesting speculation along these lines is that systems consolidation may occur as EDL in multiple stages from HC to cortex, with each faster learning region training each slower learning region (Irish & Vatansever, 2020; Remme et al., 2021; Schapiro et al., 2017).

One novel aspect of our results for memory consolidation research is that a pathway *within* the HC (EC *→* CA3) produced decontextualization. Here, we defined decontextualization as the invariance to supporting contextual input. In our initial simulations, this invariance arose to changing temporal contexts. Following findings from others that decontextualization can also arise due to invariance of learning environment (Imundo et al., 2021; S. M. Smith et al., 1978; S. M. Smith & Rothkopf, 1984), semantic context (Beheydt, 1987), and learning task (S. M. Smith & Handy, 2014, 2016), we later simulated how variable learning contexts apart from temporal drift led to similar benefits over constant learning contexts. Although to our knowledge prior paradigms have not investigated the neural locus of how decontextualization develops, a related idea called semanticization from a prominent consolidation theory (Winocur et al., 2010; Yassa & Reagh, 2013) occurs when memories lose their contextual details and retain only their central (gist-like) aspects. This theory accounts for findings that rodents with HC damage are more likely to generalize fear memories (Winocur et al., 2010), that fear memories become more generalized over time when the HC aspect of the memory trace becomes less activated (Kitamura et al., 2017; Wiltgen & Silva, 2007), and that HC amnesiac patients often appear normal when asked gist-based questions but impaired when pressed for contextual details (Nadel & Moscovitch, 1997). Therefore, semanticization is only explicitly stated to occur within the neocortex, whereas the HC is stated to support the retrieval of contextual details. Our findings showing decontextualization in HC do not oppose such a neocortical mechanism, but instead predict that such a process could initiate the formation of decontextualized traces within the HC itself (Y. Norman et al., 2021).

### Limitations

One limitation of HipSTeR is that drift-driven contextual change is simulated as a slow, passive process that is constant per unit time. However, context can also shift more rapidly (DuBrow et al., 2017), which often occurs with sudden input changes or shifts in perceived events (Antony et al., 2021; Baldassano et al., 2017; Bright et al., 2020; Brunec, Moscovitch, & Barense, 2018; Brunec et al., 2020; Clewett, Dubrow, & Davachi, 2019; Cohn-Sheehy et al., 2021; DuBrow & Davachi, 2013, 2014, 2016; Griffiths & Fuentemilla, 2020; Lu, Hasson, & Norman, 2020; Michelmann et al., 2021; Rouhani et al., 2020; Sellevoll et al., 2023; Wen & Egner, 2022; Zacks, Speer, Swallow, Braver, & Reynolds, 2007). In other words, in addition to slow drifts there are faster shifts, which allow setting up “walls” between dissimilar situational contexts nearby in time and “bridges” to similar situational contexts apart in time (Clewett et al., 2019; Cohn-Sheehy et al., 2021). Moreover, these sudden event changes can lead to more rapid forgetting of information that is more likely to rely on properly instating the temporal context (Delaney, Sahakyan, Kelley, & Zimmerman, 2010; El-Kalliny et al., 2019; Horner et al., 2016; Radvansky, Krawietz, & Tamplin, 2011), suggesting that such shifts accelerate the rate of contextual change away from the encoding context. Intriguingly, these shifts have been instantiated in sudden changes in EC (Bright et al., 2020; Tsao et al., 2018; Umbach et al., 2020) and HC activity (Griffiths & Fuentemilla, 2020), and the same EC neurons can show mixed, item-specific patterns of activity that shift with given inputs and also drift thereafter (Bright et al., 2020). Although our final environmental/pictorial decontexualization experiments begin to model such faster shifts, we hope to explore these functional activities in later versions of the model.

Another limitation is that other EC-HC subregions contribute to memory. Specifically, we have not separated LEC, which has neurons showing drifting properties (Tsao et al., 2018), from MEC. Additionally, when the HC subfields have been compared within the study in a representational drift setting, the subfield most intimately linked with changes in temporal context is CA2 (Mankin et al., 2015). CA2 and CA3 share many similarities, yet also have critical differences (Dudek, Alexander, & Farris, 2016). Here, we worked under the assumption that the EC and CA3 layers in HipSTeR combined aspects that would encompass properties of the dissociable regions. This could elucidate some aspects of the effects of CA3 in time, such as the fact that representational drift in CA1 is slower than CA3 in our model (Figure 2D) but is faster than CA3 in empirical data (Mankin et al., 2015). This could be reconciled by the fact that drift is faster in CA2 than any of the other CA regions, and these CA2 properties may be included in our CA3.

A final limitation of our model is that we did not capture some of the spacing effects produced by expanding spacing schedules. Instances in which expanding spacing is superior to equal spacing seem limited to cases in which feedback is not given after tests (Cull, Shaughnessy, & Zechmeister, 1996; Landauer & Bjork, 1978). In these training regimes, it is fairly unlikely memories will be recovered once lost, so rehearsing after an early interval after learning keeps the memory “in the running”, so to speak, whereas a slightly longer initial interval (with equal spacing) may result in a higher percentage of memories being lost and unrecoverable. In instances when feedback is given, or when restudy is used rather than retrieval, there is often no benefit of expanding spacing over equal spacing (Carpenter & DeLosh, 2005; Cull, 2000; Cull et al., 1996; Karpicke & Roediger, 2010; Küpper-Tetzel et al., 2014; Landauer & Bjork, 1978). These instances are more analogous to our model, as the target information is always available during training epochs. However, expanding performance in our model contradicted the behavioral data by being inferior to an equal schedule and not superior to a contracting schedule. There do not seem to be convincing theoretical explanations of this effect, though Küpper-Tetzel et al. (2014) have speculated that the impact of later ISIs may have a stronger influence on memory performance than earlier ISIs. Perhaps later versions of the model could attempt to employ related mechanisms.

### Conclusions and broader relevance

Given a lifetime of learning, our goal should be to optimize knowledge that can be retained over long periods of time. In this regard, the practical difficulties with conducting and publishing well-controlled research studies with long delays has likely biased the literature towards what can be assessed on relatively short time scales. Nevertheless, it is difficult to overstate the importance of spacing in long-term learning (Bahrick, 1979; Rawson & Dunlosky, 2011): even the well-known importance of testing for learning [e.g., Rowland (2014)] cannot overcome poor temporal generalization when tests are conducted at short lags, as practicing paired associates up to 10 times with one-minute lags can still produce floor (*<* 5% correct) performance a week later (Pyc & Rawson, 2009). Ultimately, a memory that remains context-dependent [e.g., Abernethy (1940); Imundo et al. (2021); Parker, Dagnall, and Coyle (2007); S. M. Smith et al. (1978); S. M. Smith and Rothkopf (1984)] may not be ideal for learning because it continues to rely on the “crutch” of context - either the presence of contextual inputs or the mental recovery of a context - for successful memory retrieval (S. M. Smith & Handy, 2014). Efforts to render memories context-independent when the context is an incidental or irrelevant aspect of the memory will ultimately benefit lifelong learning.

Recent attempts to overcome catastrophic interference and other kinds of ordering effects, such as how prior learning generalizes to new tasks, have led the artificial intelligence field to the problem of continual learning [e.g., Flesch, Balaguer, Dekker, Nili, and Summerfield (2018); Hadsell, Rao, Rusu, and Pascanu (2020); Masse et al. (2018); van de Ven, Siegelmann, and Tolias (2020)]. We hope our efforts will demonstrate the critical importance of accounting for temporal context in artificial intelligence problems. In particular, the manner in which drift between training trials - featuring different input patterns - actually improved performance in HipSTeR seems particularly important in addressing these problems.

Though representations in some regions are relatively stable over time (Dhawale et al., 2017; Katlowitz, Picardo, & Long, 2018; McGuire et al., 2022; Pérez-Ortega, Alejandre-Garćıa, & Yuste, 2021), drift occurs in many regions beyond the MTL (Deitch, Rubin, & Ziv, 2021; Driscoll, Pettit, Minderer, Chettih, & Harvey, 2017; Hyman, Ma, Balaguer-Ballester, Durstewitz, & Seamans, 2012; Margolis et al., 2012). Notably, spacing effects have been observed in numerous other learning domains, including motor-skill learning [e.g., Baddeley and Longman (1978); T. D. Lee and Genovese (1988); Shea, Lai, Black, and Park (2000)], the acquisition of math skills [e.g., Rohrer (2009)], reading [e.g., Krug, Davis, and Glover (1990)], classical conditioning [e.g., Rohrer (2009)] and extinction [e.g., Bernal-Gamboa, Gámez, and Nieto (2018); Rowe and Craske (1998)], and similar EDL mechanisms may be at play in the systems underlying these skills. Intriguingly, computational models have suggested multi-tiered learning in other domains like motor learning (Murray & Escola, 2020), suggesting parallel mechanisms in strengthening increasingly long-lasting memory traces in these systems. Additionally, there are known biological ramifications of spacing, such that spaced rather than massed inductions of long-term potentiation (LTP) result in a slower decline in LTP-induced changes (Jiang et al., 2016; Josselyn et al., 2001; Kramár et al., 2012; Scharf et al., 2002; Smolen et al., 2016). Differences in these properties across brain regions may play a critical role in how spacing and drift affect learning.

Finally, our investigations have implications for those with impairments within the EC-HC system. Anterolateral EC hypoactivity occurs in healthy aging and has been linked with memory deficits (Reagh et al., 2018), and anterolateral EC is one of the first regions to be affected by Alzheimer’s Disease [e.g., Braak and Braak (1995)]. Other long-term memory abnormalities linked with temporal lobe impairments, including accelerated long-term forgetting, which involves rapid forgetting after initially intact long-term memories (*>* 30 minutes) (Elliott et al., 2014), could ultimately rely on mechanisms uncovered here.

## Appendix

Here we provide many important parameters for understanding the HipSTeR model architecture and critical aspects about our inputs. See table captions for detailed definitions and descriptions. Please note that the full documentation and model will be included upon publication at https://github.com/ccnlab/hipster.

### Network Size Parameters

**Table 1:** Parameters for network sizes. The numbers for pool sizes indicate the number of neurons in each specific pool.

| **Network Layer** | **Neurons** |
| --- | --- |
| Input Pool Size | 7x7 |
| Input Number of Pools | 2x8 |
| ECin Pool Size | 7x7 |
| ECin Number of Pools | 2x8 |
| ECout Pool Size | 7x7 |
| ECout Number of Pools | 2x8 |
| DG Size | 60x60 |
| CA3 Size | 40x40 |
| CA1 Pool Size | 20x20 |
| CA1 Number of Pools | 2x8 |

### Training/testing input diagram

**Table 2:** Training/testing pools. Each cell represents a pool in the Input layer. TC = temporal context, B = target, A = cue, empty = no input, OC = other context for decontextualization experiments. All spacing effect simulations used TC7/TC8, whereas the decontextualization experiment used OC1/OC2. All training used B1-4 and all testing instead used Empty pools.

|  |  |
| --- | --- |
| TC7/OC1 | TC8/OC1 |
| TC5 | TC6 |
| TC3 | TC4 |
| TC1 | TC2 |
| B3/Empty | B4/Empty |
| B1/Empty | B2/Empty |
| A3 | A4 |
| A1 | A2 |

### Parameter changes from default model

**Table 3:** Model parameters that diverged from the default (Zheng et al., 2022). These changes were made to account for different network sizes, specifically the change from 2×3 input pools used in Zheng et al. (2022) to the 2×8 pools here.

| Area | Param | Value | Default |
| --- | --- | --- | --- |
| ECout | ECoutToECin.WtScale.Rel | 0.4 | 0.5 |
| DG | Inhib.Layer.Gi | 2.95 | 3.8 |

*

